# A Novel DNA Chromatography Method to Distinguish *M. abscessus* Subspecies and Macrolide Susceptibility

**DOI:** 10.1101/2020.09.17.292417

**Authors:** Mitsunori Yoshida, Sotaro Sano, Jung-Yien Chien, Hanako Fukano, Masato Suzuki, Takanori Asakura, Kozo Morimoto, Yoshiro Murase, Shigehiko Miyamoto, Atsuyuki Kurashima, Naoki Hasegawa, Po-Ren Hsueh, Satoshi Mitarai, Manabu Ato, Yoshihiko Hoshino

## Abstract

**Rationale:** The clinical impact of infection with *Mycobacterium abscessus* complex (MABC), a group of emerging non-tuberculosis mycobacteria (NTM), is increasing. *Mycobacterium abscessus* subsp. *abscessus*/*bolletii* frequently shows natural resistance to macrolide antibiotics, whereas *Mycobacterium abscessus* subsp. *massiliense* is generally susceptible. Therefore, rapid and accurate discrimination of macrolide-susceptible MABC subgroups is required for effective clinical decisions about macrolide treatments for MABC infection.

**Objectives:** To develop a simple and rapid diagnostic that can identify MABC isolates showing macrolide susceptibility.

**Methods:** Whole genome sequencing (WGS) was performed for 148 clinical or environmental MABC isolates from Japan to identify genetic markers that can discriminate three MABC subspecies and the macrolide-susceptible *erm*(41) T28C sequevar. Using the identified genetic markers, we established PCR based- or DNA chromatography-based assays. Validation testing was performed using MABC isolates from Taiwan.

**Measurements and Main Results:** We identified unique sequence regions that could be used to differentiate the three subspecies. Our WGS-based phylogenetic analysis indicated that *M. abscessus* carrying the macrolide-susceptible *erm*(41) T28C sequevar were tightly clustered, and identified 11 genes that were significantly associated with the lineage for use as genetic markers. To detect these genetic markers and the *erm*(41) locus, we developed a DNA chromatography method that identified three subspecies, the *erm*(41) T28C sequevar and intact *erm*(41) for MABC in a single assay within one hour. The agreement rate between the DNA chromatography-based and WGS-based identification was 99.7%.

**Conclusions:** We developed a novel, rapid and simple DNA chromatography method for identification of MABC macrolide susceptibility with high accuracy.

## Introduction

The *Mycobacterium abscessus* (heterotypic synonym; *Mycobacteroides abscessus*)(1, 2) complex (MABC) is a group of rapid-growing non-tuberculosis mycobacteria (NTM) that includes three subspecies: *M. abscessus* subsp. *abscessus* (*M. abscessus*), *M. abscessus* subsp. *massiliense* (*M. massiliense*), and *M. abscessus* subsp. *bolletii* (*M. bolletii*)(3, 4). MABC causes a range of clinical infections including chronic pulmonary disease even in immunocompetent persons, as well as postsurgical or traumatic infections and skin and soft tissue infections(4–8).

Among NTM infections, treatment outcomes for MABC infections are relatively worse and the in-hospital mortality rate can reach 16%(5, 9–13). These poor outcomes are due in part to the extensive antibiotic resistance of MABC(14). However, some MABC patients achieve good clinical outcomes with standard antibiotic regimens(15–17). Out of the three subspecies, *M. massiliense* is susceptible to macrolide antibiotics whereas *M. abscessus* and *M. bolletii* are resistant(18, 19). In the presence of macrolide antibiotics *M. abscessus* and *M. bolletii* exhibit inducible expression of erythromycin ribosomal methylase (*erm*)(41), which produces the Erm protein that reduces macrolide affinity for the ribosome exit tunnel(20–22). *M. massiliense* harbors a truncated *erm*(41) that produces inactive Erm(41). A T-to-C sequence variant (sequevar) at position 28 (T28C) of the *erm*(41) gene also results in production of an inactive enzyme, and does not result in inducible macrolide resistance of *M. abscessus* and *M. bolletii*, which are generally macrolide resistant. These observations indicate that determination of subspecies and detection of intact *erm*(41) and the *erm*(41) T28C sequevar of MABC can inform prediction of clinical course and treatment outcome. In fact, the 2020 ATS/ERS/ESCMID/IDSA Clinical Practice Guideline strongly recommends a macrolide-containing multidrug treatment regimen for patients with MABC respiratory disease caused by strains without inducible macrolide resistance(23). Accordingly, discrimination of subspecies and identification of the *erm*(41) T28C sequevar is crucial.

Sequencing of single 16S rRNA or the RNA polymerase beta subunit (*rpoB*) cannot distinguish MABC subspecies because these loci are nearly identical(13, 24). Several studies have examined use of multi-locus sequencing typing (MLST) of housekeeping genes to separate subspecies(25–28). Advances in whole genome sequencing (WGS) technology allowed MABC clinical isolates to be phylogenetically divided into three subspecies even at the whole genome level(29, 30). However, detection of the *erm*(41) T28C sequevar still requires sequencing of the entire *erm*(41) gene. Although MLST and/or WGS analyses allow discrimination of *erm* genes, these analyses are time-consuming and labor intensive in clinical practice. Thus, novel assays that are simple and rapid yet retain discriminatory power required to distinguish subspecies and to identify the *erm*(41) T28C sequevar are needed.

We previously reported a PCR-based method to differentiate MABC subspecies, but the capacity of this test was limited(31). In the present study, we analyzed WGS data for 148 MABC isolates from Japan to explore genetic markers associated with each subspecies and the *erm*(41) T28C sequevar. We propose a novel, rapid and easy-to-use DNA chromatography method that can identify all MABC subspecies as well as intact *erm*(41) and the *erm*(41) T28C sequevar in a single assay.

## Materials and methods

For further details on the applied methods, *see* the online data supplement

### Bacterial isolates

A total of 147 MABC clinical isolates and one environmental isolate (strain MabLRCB1) obtained for differential diagnosis at 19 hospitals (listed in Acknowledgments) in Japan were considered. Of the clinical isolates, 138 originated in the respiratory system, 8 were isolated from skin lesions, and 1 strain was isolated from a blood sample (Table S1). Another 103 clinical isolates were obtained from Taiwan National University Hospital (Yoshida *et al.*, manuscript in preparation, Table S2). All strains were classified as MABC using a DDH Mycobacteria kit (Kyokuto Pharmaceutical Industrial, Tokyo, Japan) or by MALDI-TOF MS (Bruker Daltonics, Billerica, MA, USA). *M. abscessus* subsp. *Abscessus* JCM 13569^T^ (ATCC 19977), *M. abscessus* subsp. *massiliense* JCM 15300^T^ and *M. abscessus* subsp. *bolletii* JCM 15297^T^ (BD) type strains were obtained from the Japan Collection of Microorganisms of the Riken Bio-Resource Center (BRC-JCM; Ibaraki, Japan). All bacterial strains/isolates were subcultured on 2% Ogawa egg slants or 7H10 agar plates supplemented with 10% OADC.

### PCR assays for discriminating *M. abscessus* subspecies and *erm*(41) T28C sequevar

Single-PCR and multiplex PCR assays differentiating *M. abscessus*, *M. massiliense*, and *M. bolletii*, as well as the *erm*(41) T28C sequevar were conducted essentially as described previously(31) using newly constructed primers (*see* supplemental methods). Briefly, template DNA for PCR assays was isolated from one loopful of a mycobacterial colony grown on 7H10 medium that were resuspended in 300 μl sterilized water, boiled at 95 °C for 15 min and frozen at -30 °C. PCR amplification was performed using a Mastercycler gradient (Eppendorf) with 95 °C for 10 min; 30 cycles of 95 °C for 30 sec, 60 °C for 30 sec, and 72 °C for 40 sec; and extension at 72 °C for 10 min. Amplification to identify the *erm*(41) T28C sequevar was performed in the Mastercycler gradient using 95 °C for 10 min; 35 cycles of 95 °C for 1 min, 62 °C for 1 min, and 72 °C for 1 min; and extension at 72 °C for 4 min. The PCR products were separated by 2% agarose gel electrophoresis and stained with ethidium bromide. The analytical limit of detection of the multiplex PCR assay was estimated by applying serial dilutions of DNA from *M. abscessus* ATCC19977, *M. massiliense* JCM 15300, and *M. bolletii* BD, in addition to several clinical MABC isolates. To assess the multiplex PCR assay specificity, several laboratory and clinical isolates were used including the *M. avium* complex (10 clinical isolates and one laboratory stain), *M. conceptionense, M. fortuitum, M. gordonae, M. houstonense, M. kansasii, M. leprae* (3 clinical isolates and one laboratory stain)*, M. lentiflavum, M. peregrinum, M. salmoniphilium*, *M. senegalense*, *M. shimoidei, M. smegmatis*, *M. szulgai, M. triplex, M. tuberculosis* (10 clinical isolates and one laboratory stain), and *M. xenopi*.

### DNA chromatography assay for discriminating MABC subspecies, intact *erm*(41) and the *erm*(41) T28C sequevar

We applied a DNA chromatography method that was described elsewhere(32–34) to distinguish the MABC subspecies, intact *erm*(41) gene and the *erm*(41) T28C sequevar. This assay comprises PCR amplification and amplicon detection. All primers used in the assay are listed in Table 3. Briefly, primers with 5’ tags, which have a domain that anneals to the target sequence and a tag domain that hybridizes to a single-stranded DNA probe on the chip or gold nanoparticle, were used for amplification. DNA extraction was performed essentially as described previously(6, 35). Total genomic DNA was extracted from frozen samples (as described above) using a Kaneka easy DNA extraction kit for *Mycobacteria* (KANEKA, Osaka, Japan). PCR was performed using a 20 μl mixture containing 10 μl PCR Mix (KANEKA, Osaka, Japan), 5 μl primer mix (5 primer sets, 0.5 μM each, Table 3), 1 μl template DNA. Amplification was performed in a Life ECO thermocycler (BIOER Co. Ltd., Hangzhou, China) with 25 °C for 5 min and 94 °C for 1 min, followed by 35 cycles of 94 °C for 5 sec, 65 °C for 10 sec, and 72 °C for 15 sec. An aliquot of the amplicons supplemented with 70 μl development buffer (KANEKA, Osaka, Japan) was applied to the detection strip sample pad (KANEKA, Osaka, Japan). After 10 min, blue lines were confirmed visually. Sensitivity and specificity tests for the DNA chromatography were performed as described above.

## Results

### Subspecies identification based on core gene alignment of MABC isolates

In the context of genome-based taxonomy, phylogeny involving concatenated core gene alignments is frequently used instead of 16S rRNA or *rpoB* gene sequences. We first examined whether a concatenated sequence of core genes, which are defined as homologous genes present in all strains examined, could distinguish the three MABC subspecies. Using a concatenated sequence of 2,957 core genes, the 148 MABC isolates could clearly be divided into three clades, with 92 (62.2%), 52 (35.1%), and 4 (2.7%) isolates identified as *M. abscessus*, *M. massiliense*, and *M. bolletii*, respectively (Fig. 1). This result is consistent with our previous report showing that multi-locus sequence typing (MLST) of *rpoB*, *hsp65*, and the ITS region could discriminate the three subspecies with 97.5% accuracy(31). Moreover, most isolates could be phylogenetically categorized in agreement with the core-locus phylogeny, although 4 (2.7%) were inconsistently categorized (Fig. S1A). Another group proposed a MLST scheme using seven housekeeping genes(28). We confirmed that this scheme could reliably discriminate the three MABC subspecies (Fig. S1B). We also confirmed the subspecies identification by calculating the average nucleotide identity (ANI) for all MABC isolates (Fig. S2). The minimum ANI within each of the three subspecies was 98.4 (within *M. abscessus*), 98.3 (within *M. massiliense*), and 99.1 (within *M. bolletii*), while the maximum ANI between subspecies was 97.5 (between *M. abscessus* and *M. massiliense*), 97.7 (between *M. abscessus* and *M. bolletii*), and 97.0 (between *M. massiliense* and *M. bolletii*). These results indicated that phylogenetic analysis based on core-locus alignment indeed differentiated the three MABC subspecies and suggested that their subspecies boundaries were approximately 98% ANI.

**Figure 1.**
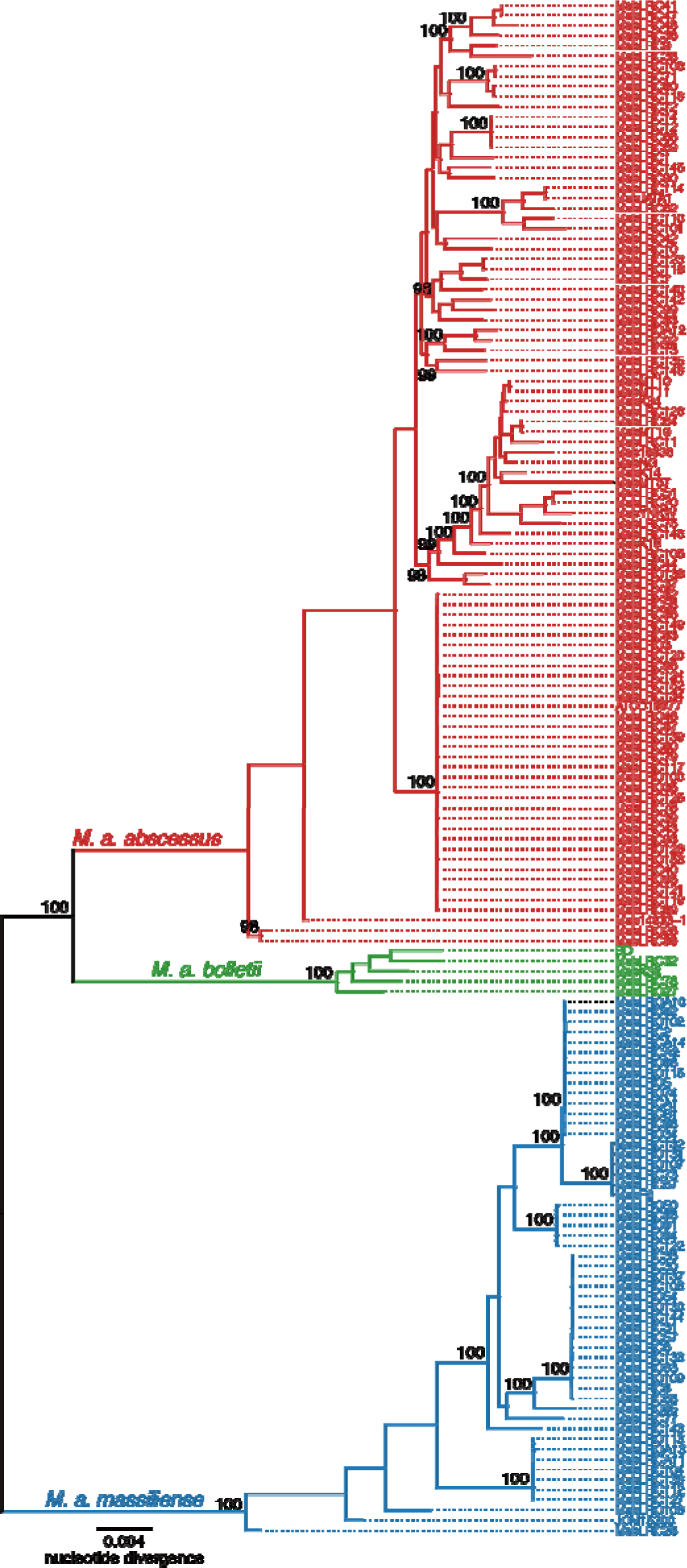
Maximum likelihood core-gene phylogeny of 148 clinical and environmental isolates of MABC. Core genome alignment of 148 isolates and three reference strains (*M. abscessus* ATCC19977, *M. massiliense* JCM 15300, and *M. bolletii* BD) of MABC was generated by Roary(51). An alignment containing 62,196 variable positions was used with RAxML to construct a maximum likelihood tree(52) having 300 bootstrap replicates. Bootstrap values > 90% for the major nodes are shown. Scale bar indicates the mean number of nucleotide substitutions per site (SNP/site) on the respective branch. Samples are highlighted based on inclusion in three major clusters corresponding to MABC subspecies.

### Multiplex PCR assay for discriminating the three MABC subspecies

Since WGS and/or MLST are not feasible for clinical settings, we sought to develop an alternative method to distinguish the three subspecies of MABC. We focused on “genetic markers” specific to each subspecies. We first aligned the complete genome sequences of type strains (ATCC 19977, JCM 15300, and BD) and draft genome sequences of 14 representative clinical isolates with progressiveMauve(36) and visually identified unique insertion/deletion (indel) regions in each of the three subspecies. We then designed three primer sets specific for sequences around the indel regions for size-based differentiation of PCR amplicons (Fig. 2A, Table 1). All primer sets could sharply discriminate reference strains and clinical isolates as evidenced by a single band on an agarose gel (Fig. 2B and Fig. S3). A sensitivity test showed the limit of detection was 100 pg DNA (Fig. S4). All other laboratory and clinical isolates of NTM and *M. tuberculosis* tested were negative in this PCR assay (Fig. S5). We subsequently examined the primer set accuracy using the 148 MABC isolates, and confirmed that 91/92 *M. abscessus* (98.9%), 52/52 *M. massiliense* (100%), and 4/4 *M. bolletii* (100%) isolates were in agreement with the WGS-based subspecies identification, and the overall agreement rate between the PCR assay and WGS-based subspecies identification was 99.3% (95% CI: 96.3% to 100%). Results of our previous MLST(31) showed several sequence variants in the isolates, but the PCR assay could still differentiate between strains (Fig. 2B, Fig. S3). Notably, discordant sequencing type 4 (MabLRC28 and MabLRC86) and ds type 5 (MabLRC28) were distinguished as *M. abscessus* and *M. massiliense*, respectively, in accordance with WGS-based subspecies identification (Fig. 2). Although our previous multiplex PCR assay(31) could not distinguish all *M. bolletii* from others (Table S1), the present PCR assay could distinguish them. Moreover, the agreement rate with WGS-based subspecies identification was significantly higher (*P* < 0.01, two-proportion *Z* test) than that of the previous multiplex PCR assay (92.6%, 95% CI: 87.1% to 96.2%).

**Figure 2.**
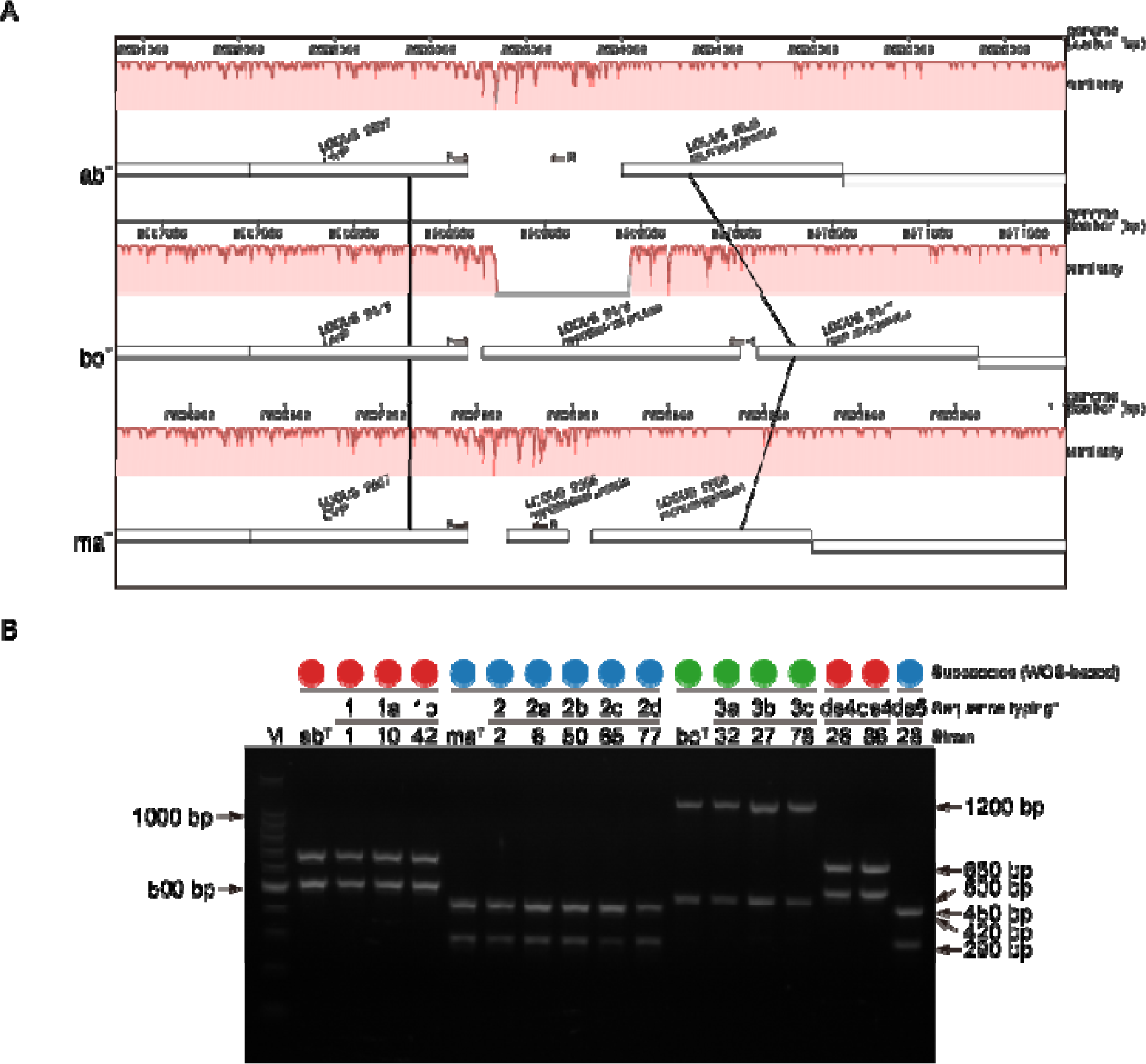
A. Example of indels among MABC subspecies. A progressiveMauve alignment of the three reference strains of MABC is shown. Each genome is laid out in a horizontal track and white boxes indicate coding sequences annotated by dfast_core(53). A colored similarity plot is shown for each genome; the height is proportional to the sequence identity in that region. F and R indicate primer position of MAB2613F and MAB2613R (listed in Table 1), respectively, in each reference strain genome. **B. Representative multiplex PCR results for reference strains and clinical isolates amplified with primer pair MAB2613F and MAB2613R and primer pair MAB_1655F and MAB_1655R.** Types 1, 1a, 1b, 2, 2a, 2b, 2c, 2d, 3, 3a, 3b, 3c, ds4 and ds5 are the sub-groupings (sequevars) of the clinical isolates based on their sequences [Table 3 of the previous article(31)]. Numerals below the sequevars correspond to reference strains (ab^T^, ma^T^, or bo^T^) or the strain numbers of clinical isolates described in the previous article(31). Colored circles correspond to each member of MABC determined by WGS-based analysis. Lanes: M, DNA marker (100 bp ladder); ab^T^, *M. abscessus* (ATCC19977); ma, *M. massiliense* (JCM15300); bo, *M. bolletii* (BD). The PCR reaction products were electrophoresed on 2% agarose gels.

**Table 1.**
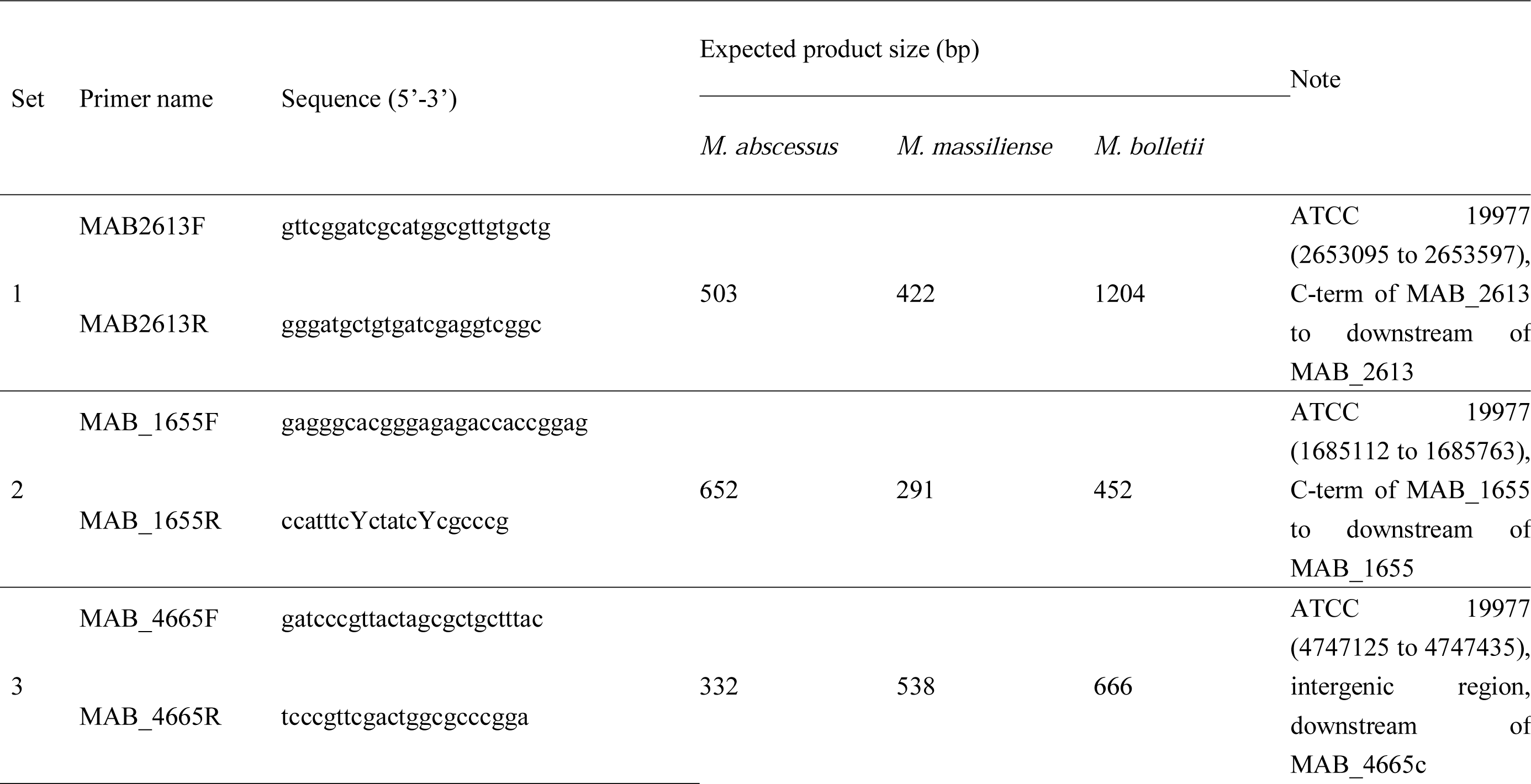
Primers for discrimination of MABC subspecies.

### Multiplex PCR assay for discriminating the *erm*(41) T28C sequevar of MABC

MABC have an Erm(41)-mediated inducible mechanism for resistance to macrolide antibiotics. Earlier studies demonstrated that a T-to-C substitution at position 28 in the *erm* gene results in macrolide antibiotic susceptibility(20, 37). Thus, discriminating isolates that carry the *erm*(41) T28C sequevar could guide antibiotic selection. To identify genetic markers associated with the *erm*(41) T28C sequevar, we investigated the *erm*(41) genotype of the 148 MABC isolates and their phylogenetic relationship (Fig. 3). Of the 92 *M. abscesssus*, 17 (18.5%) had the *erm*(41) T28C mutation and all but MabLRC70 were susceptible to clarithromycin (CAM) (Table S1). Notably, our phylogenetic analysis showed that these clinical isolates were tightly clustered (Fig. 3). Scoary analysis of lineage-associated genes indicated that 68 genes were significantly associated with the lineage to which all *M. abscessus* with the *erm*(41) T28C sequevar belonged (Bonferroni corrected *P*-value < 1E-8, sensitivity >80% and specificity > 80%, Table S4). In a whole-genome alignment, 11/68 lineage-associated genes having the highest sensitivity and specificity were on a lineage-specific genetic locus (Fig. 4A). To determine whether these findings applied to other sample sets, we used public WGS data for MABC clinical isolates from two European countries(38, 39). A phylogenetic analysis based on the core-gene alignment indicated that the *M. abscessus erm*(41) T28C sequevar was also clustered and had the abovementioned genetic locus (Fig. S6). Among primer sets designed to amplify part of the genetic locus (Table 2, Fig. 4A), one set detected only the *M. abscessus erm*(41) T28C sequevar with a single band (Fig. 4B). Among the 148 MABC isolates, all 17 *M. abscessus erm*(41) T28C sequevars were positive in the PCR assay, whereas all other clinical isolates examined, except for MabMT19 and MabLRC77, were negative (Table S1). The agreement between the multiplex PCR assay results with WGS-based discrimination of the *erm*(41) T28C sequevar was 98.6% (95% CI: 95.2% to 99.8%).

**Figure 3.**
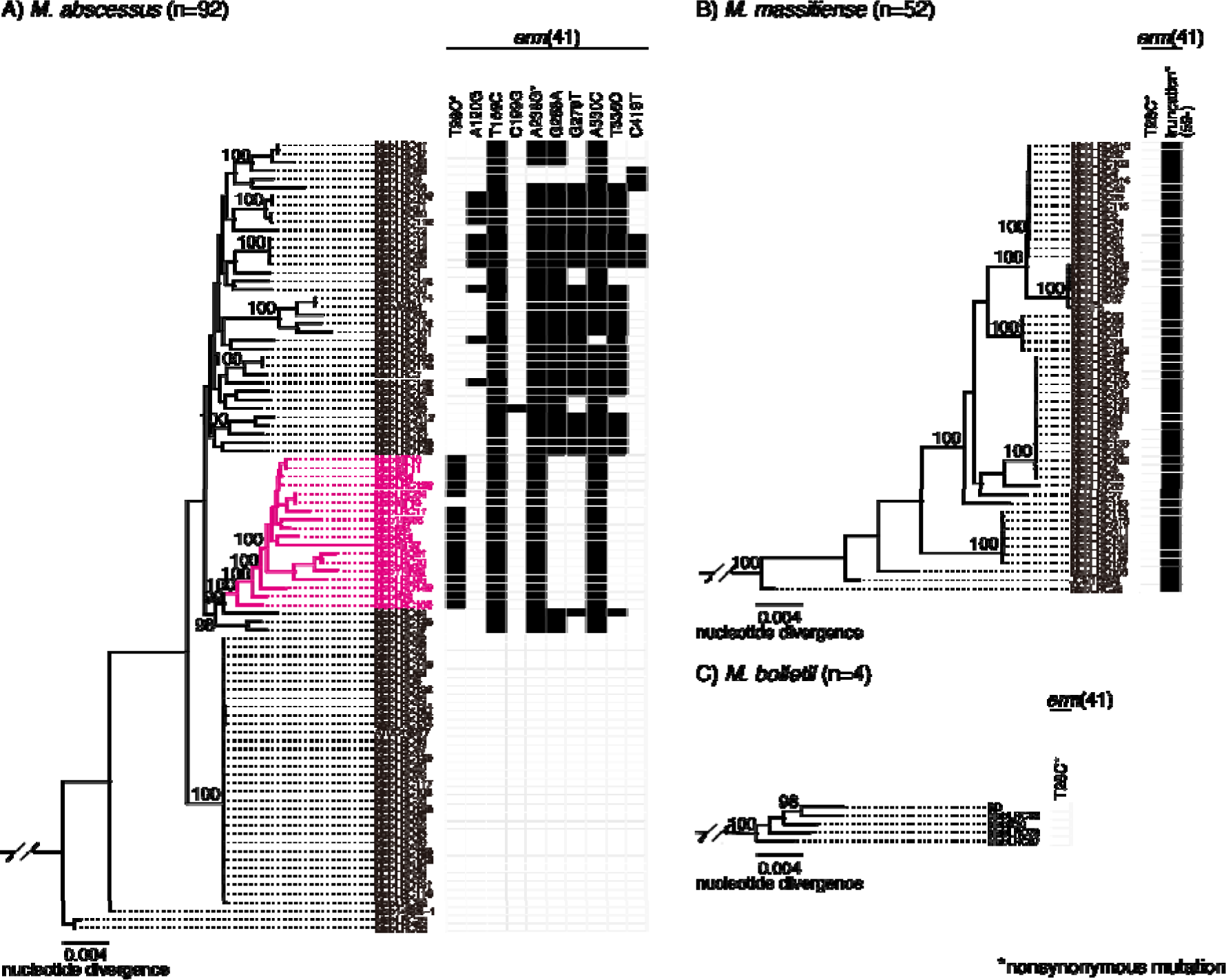
Macrolide susceptibility-associated genotypes of 148 MABC isolates. Maximum likelihood core-gene phylogeny of A) *M. abscessus*, B) *M. massisliense*, C) *M. bolletii* correspond to those depicted in Figure 1. The presence (black) and absence (gray) of macrolide resistance-associated mutations is indicated. The presence of a T-to-C substitution in position 28 or a truncation of the *erm*(41) gene, which are both associated with inducible resistance to macrolides, was detected. Substitutions or truncations with asterisks indicate non-synonymous mutations. The lineage to which all *M. abscessus erm*(41) T28C mutants belong is highlighted in magenta. The maximum likelihood trees, bootstrap values and scale bars correspond to those depicted Figure 1.

**Figure 4.**
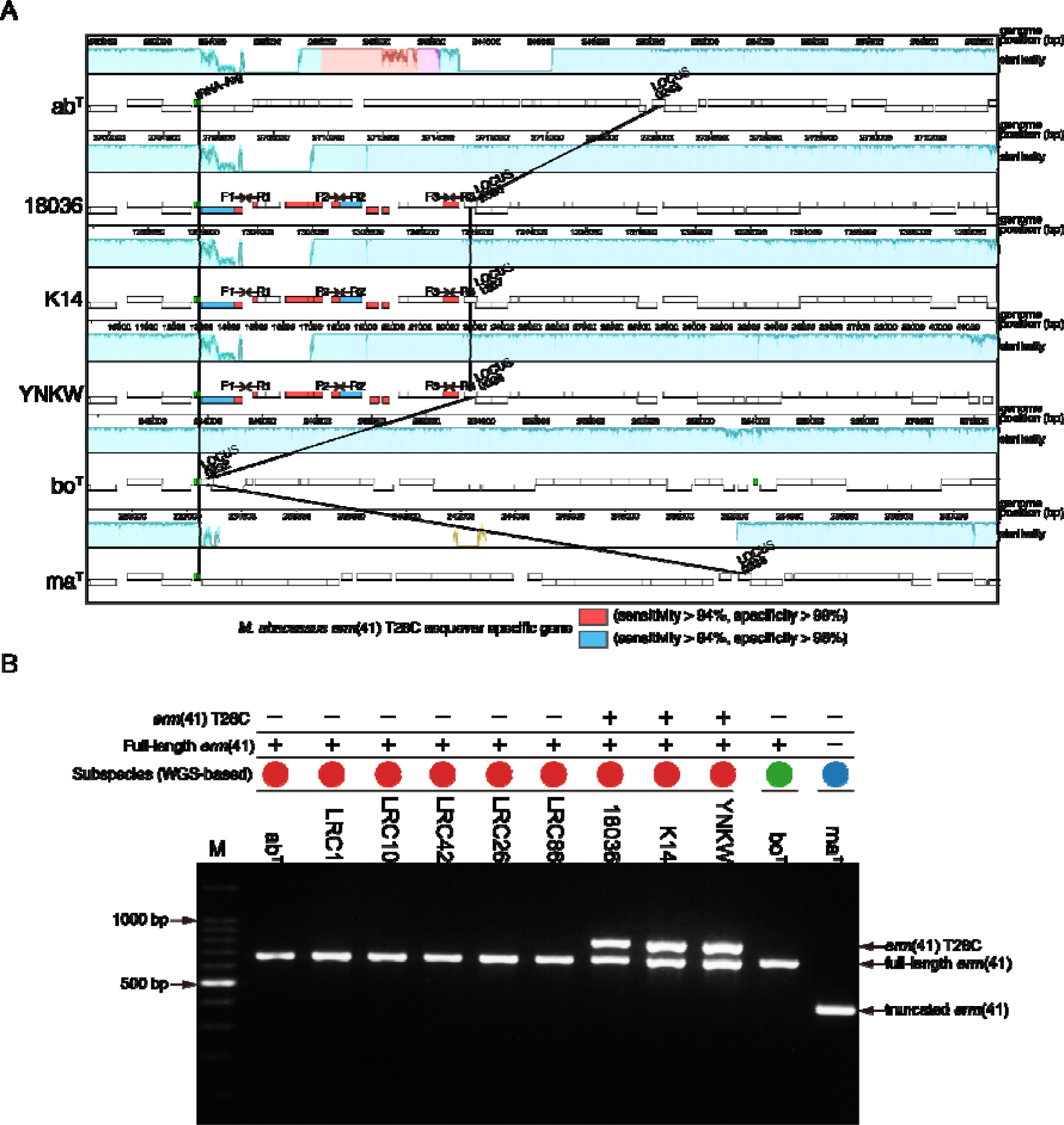
A. Visualization of lineage-specific genomic loci. A progressiveMauve alignment of three clinical isolates carrying the *erm*(41) T28C mutation and the reference strains is shown. Boxes indicate coding sequences annotated by dfast_core(53). Red and blue boxes indicate genes that are significantly associated with the lineage to which all *M. abscessus erm*(41) T28C mutants belong (see Methods and Table S3). F1, R1, F2, R2, F3, and R3 indicate primer position of MAB18036_2551F, MAB18036_2551R, MAB18036_2558F, MAB18036_2558R, MAB18036_2560F, and MAB18036_2560R (listed in Table 2), respectively, in each genome of the presented clinical isolates. A similarity plot for each genome is colored as described for Figure 2A. **B. Representative multiplex PCR results to identify macrolide susceptibility in MABC**. PCR was performed for the reference strains and clinical isolates were amplified with primer pairs MAB18036_2558F and MAB18036_2558R (listed in Table 2) and primer pair ermF (gaccggggccttcttcgtgatc) and ermR (agcttccccgcaccgattcca)(54). Colored circles correspond to each member of MABC determined by WGS-based analyses: red, *M. abscessus*; blue, *M. massiliense*; green, *M. bolletii*. The presence (+) or absence (-) of genotypes associated with the inducible macrolide resistance are shown. Lanes: M, DNA marker (100 bp ladder); ab^T^, *M. abscessus* (ATCC19977); ma^T^, *M. massiliense* (JCM15300); bo^T^, *M. bolletii* (BD). The PCR reaction products were electrophoresed on 2% agarose gels.

**Table 2.**
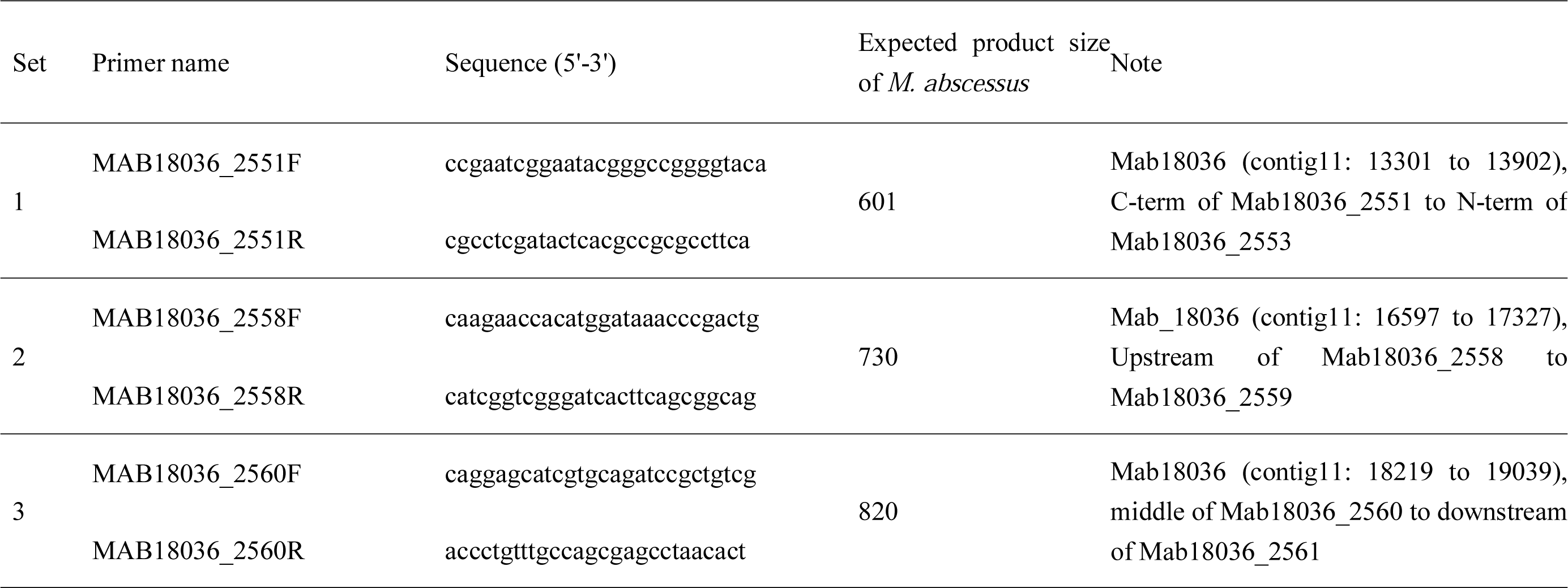
Primers for discrimination of the erm(41) T28C sequevar.

**Table 3.**
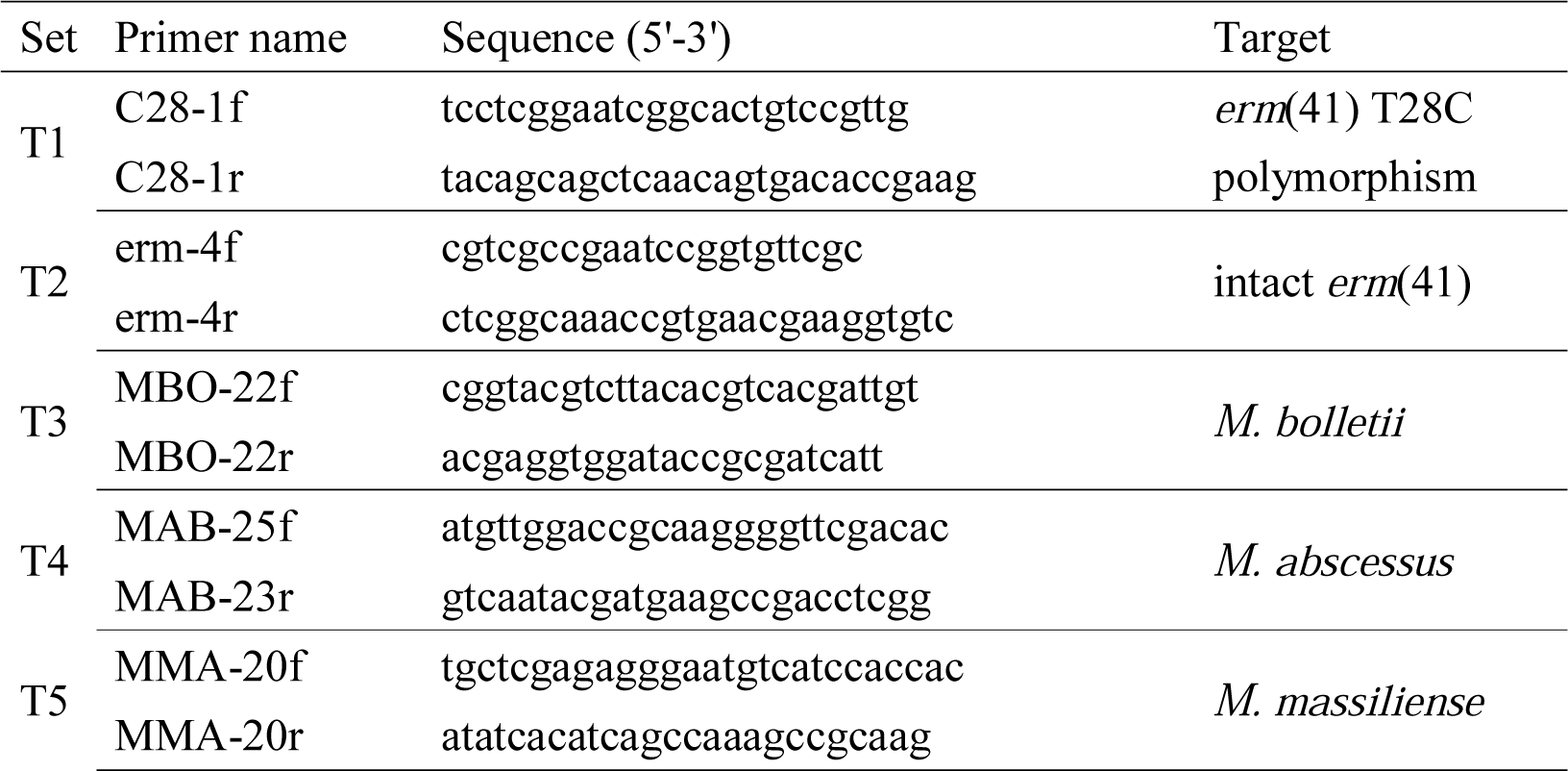
DNA chromatography primers.

### DNA chromatography to discriminate subspecies and macrolide susceptibility of MABC

Based on the results for the two PCR-based assays to discriminate MABC subspecies and the *erm*(41) T28C sequevar, we developed a simple DNA chromatography-based assay to discriminate MABC subspecies, intact *erm*(41), and the *erm*(41) T28C sequevar. We could discriminate subspecies and macrolide resistance in a single assay that in a sensitivity test had a detection limit of 10 pg DNA (Fig. 5), which was more sensitive than that of the multiplex PCR assay (Fig. S4, S7). All other laboratory and clinical isolates of NTM as well as *M. leprae* and *M. tuberculosis* that were tested were negative in the assay (Fig. S8 and data not shown). Using the 148 MABC isolates, we also examined the agreement between the DNA chromatography assay and WGS-based discrimination. All *M. abscessus* (n=92), *M. massiliense* (n=52), and *M. bolletii* (n=4) isolates were positive with T4, T5, and T3 bands respectively, while all other isolates identified as the remaining two subspecies were negative for these bands (Table 4, Table S1). All *M. abscessus* carrying the *erm*(41) T28C sequevar were positive, whereas other strains carrying wild-type T28 were negative for the T1 band, except for MabMT19 and MabLRC77 clinical isolates. All *M. abscessus* (n=92) and *M. bolletii* (n=4) strains carrying a intact *erm*(41) gene were positive for the T2 band but all *M. massiliense* (n=53) carrying a truncated *erm*(41) gene were negative, which is consistent with WGS-based analyses (Table 4, Fig. 3). Overall agreement between the DNA chromatography results with WGS-based discrimination was 99.7% (95% CI: 99.0% to 100%). We also used the DNA chromatography method to analyze another sample set comprising 103 MABC clinical isolates from Taiwan for validation (Table S2). Using this method, within only a few hours we could determine that 49, 2, and 50 clinical isolates were *M. abscessus*, *M. bolletii* and *M. massiliense*, respectively; subspecies of TJMA-002 and TJMA-104 (1.9%) were not determined because these isolates showed multiple bands in subspecies identification (Table S2). Of 47 clinical isolates showing the T4 band, 12 also showed the T1 and T2 band, which corresponded to the *M. abscessus erm*(41) T28C sequevar, while 37 isolates showed only the T2 band corresponding to *M. abscessus* with an intact *erm*(41) gene. Two clinical isolates showing the T3 band also showed the T2 band, indicating that all *M. bolletii* had an intact *erm*(41) gene. Of 50 isolates showing the T5 band, 47 did not show the T2 band, indicating an *erm*(41) gene truncation. However, the remaining three isolates (TJMA-024, TJMA-041, TJMA-046) unexpectedly showed the T2 band. Using PCR amplification, TJMA-024 and TJMA-041 isolates had intact *erm*(41), whereas TJMA-046 had both an intact and truncated *erm*(41) gene (data not shown). These results suggested that TJMA-024 and TJMA-041 were *M. massiliense* with an intact *erm*(41) gene and TJMA-041 was probably a mixed isolate of *M. abscessus* or *M. bolletii* having an intact *erm*(41) gene and *M. massiliense* with a truncated *erm*(41) gene.

**Figure 5.**
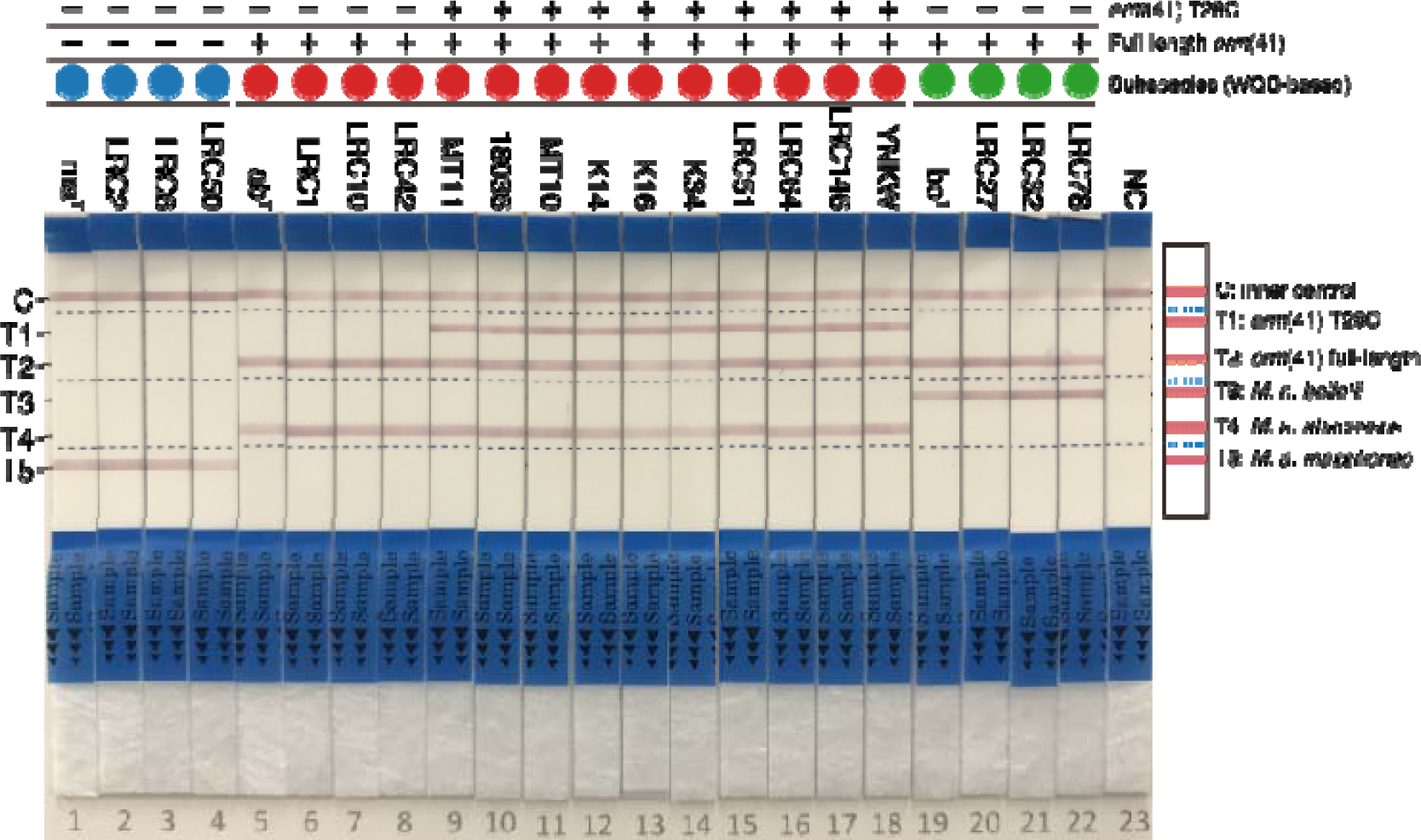
DNA chromatography to differentiate subspecies and macrolide susceptibility of MABC. DNA chromatography results for reference strains (abT, maT, or boT) or representative clinical isolates are shown. Colored circles above the strain names correspond to each member of MABC determined by WGS-based analyses: red, *M. abscessus*; blue, *M. massiliense*; green, *M. bolletii*, respectively. The presence (+) or absence (-) of genotypes associated with inducible macrolide resistance are shown. Bands: C, inner (negative) control; T1, *erm*(41) T28C polymorphism; T2, intact *erm*(41) genes; T3, *M. bolletii*; T4, *M. abscessus*; T5, *M. massiliense*.

**Table 4.**
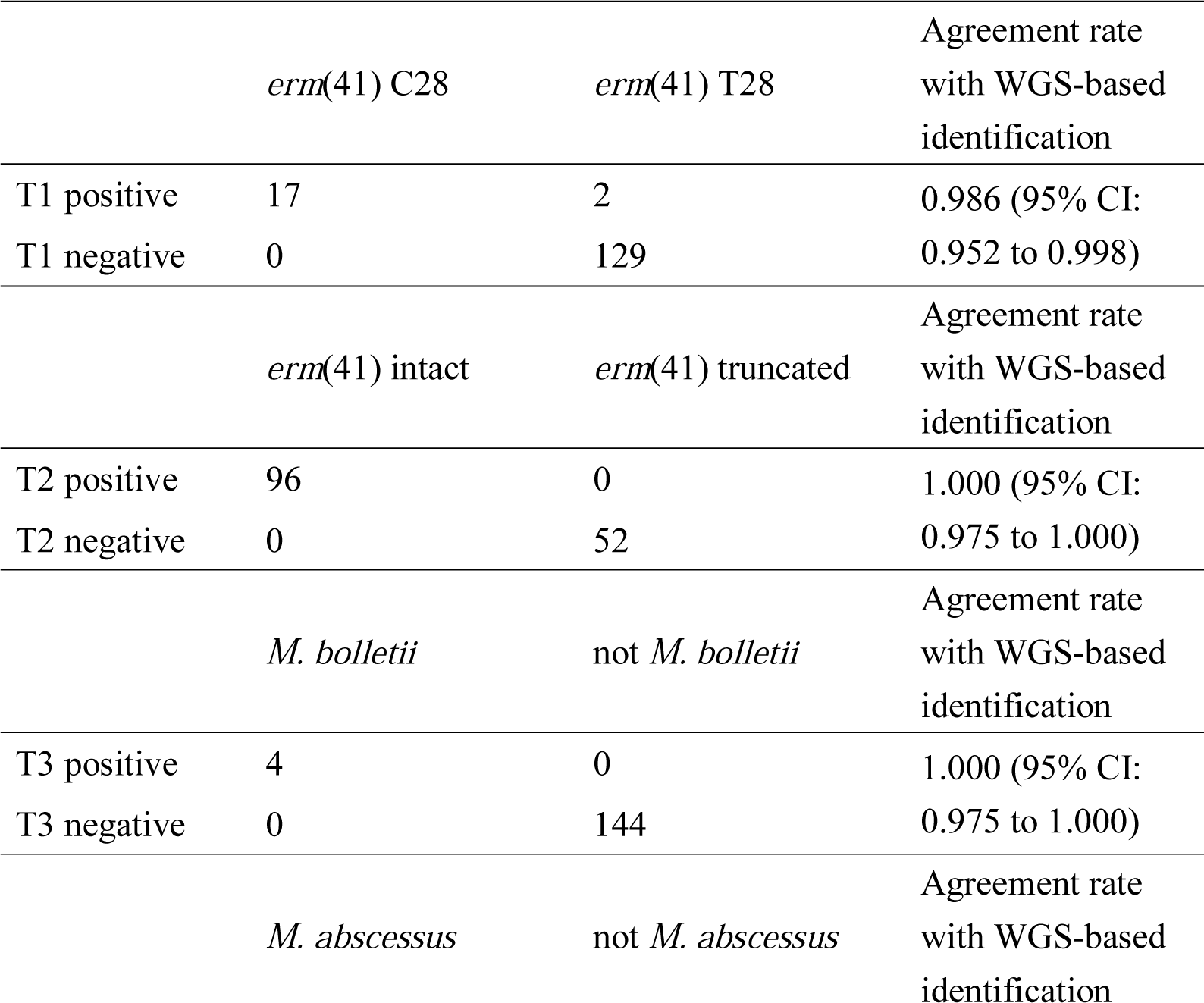
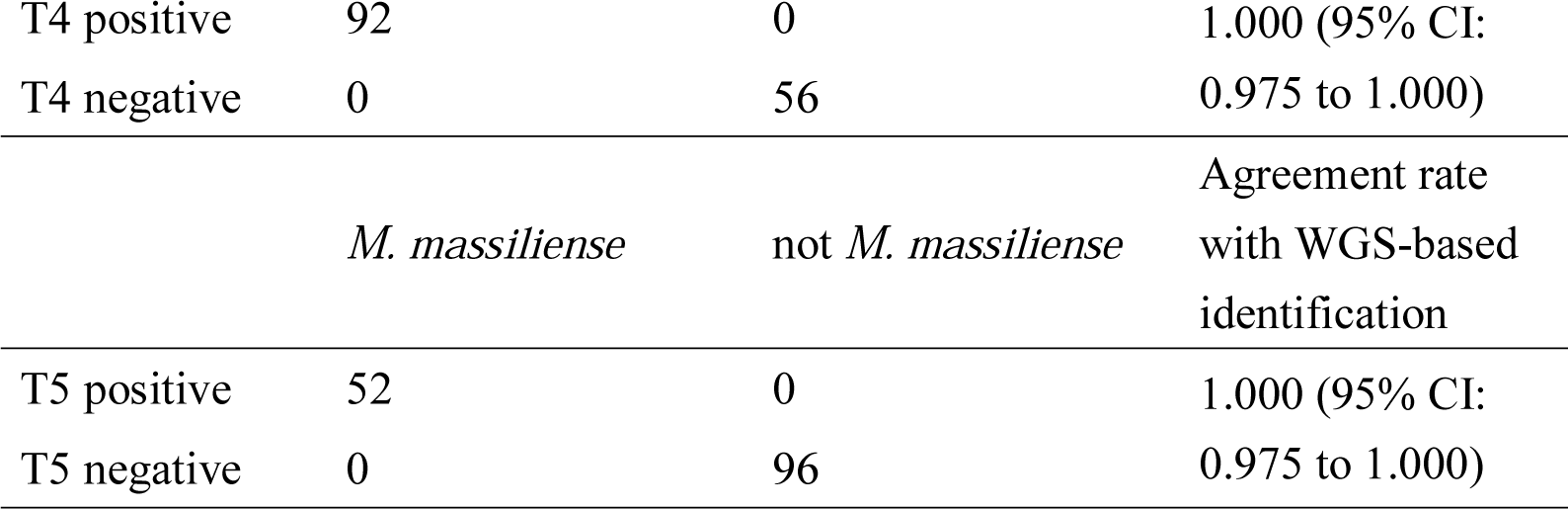
Accuracy of DNA chromatography test using MABC isolates from Japan (n = 148).

## Discussion

MABC is the most frequent clinical isolate of rapidly growing mycobacteria and an increased emergence has recently been observed in Japan and other developed countries(40–42). Commercially available DNA-DNA hybridization kits or MALDI-TOF MS are available in clinical laboratories in Japan and Taiwan to identify mycobacterium isolates(43, 44), but they cannot discriminate between either subspecies or the macrolide-susceptible *erm*(41) truncation and T28C polymorphism(45). Since macrolide susceptibility is crucial for effective treatment of MABC infection, here we developed a novel multiplex PCR and DNA chromatography method to identify subspecies and macrolide susceptibility. This assay allows rapid and accurate identification of inducible-macrolide resistance without need for sequencing of the *erm*(41) gene and/or 14-day drug susceptibility testing as recommended in the recent ATS/ERS/ESCMID/IDSA Clinical Practice Guideline(23). Our methodology is based on two findings about MABC genome architecture: (i) indel regions are robustly conserved at the subspecies level, and (ii) the macrolide-susceptible *M. abscessus* T28C sequevar is phylogenetically clustered and shares specific genetic loci. We thus used subspecies-associated and *erm*(41) T28C sequevar-associated genomic sequences as genetic markers to predict macrolide susceptibility. By combining detection of these genetic markers with simple DNA chromatography without DNA degeneration processes, our assay could discriminate both subspecies and their macrolide susceptibility more quickly and easily than previously described MLST or WGS methods(43).

DNA chromatography produces clear visual results using only a thermocycler rather than more expensive and complex genome sequencers or MALDI-TOF MS instruments. The one hour turnaround time between DNA extraction to availability of results would substantially reduce the need for MLST or WGS to differentiate MABC subspecies and inducible macrolide susceptibility. Compared with the commercially available GenoType NTM-DR kit (Hain Lifesciences GmbH, Bruker Corporation, Nehren, Germany), the present DNA chromatography method is simpler (DNA chromatography vs. Southern blotting) and faster (1 hour vs. >4 hours). Furthermore, the accuracy of subspecies identification is higher (98%-100% vs. 92%-100%), and the ability to discriminate the *erm*(41) T28C sequevar is comparable(46–48), although the GenoType NTM-DR kit can detect additional acquired macrolide resistance and amikacin resistance.

Our DNA chromatography method had an analytical limit of detection of 10 pg DNA, which is substantially more sensitive than the GenoType NTM-DR kit (2 ng)(48). This amount of DNA theoretically corresponds to approximately 2.2 x 10^3^ MABC cells (DNA content/cell = genome size(bp)/0.978×10^9^ 5.11×10^-3^)(49). There was no cross-reactivity with *M. leprae*, *M. tuberculosis*, MAC, or other representative NTM. These observations suggest that the DNA chromatography assay can identify MABC isolates directly from liquid MGIT or solid Löwenstein-Jensen (LJ) or Ogawa cultures at early time points after decontamination of non-mycobacterial organisms. However, we have not yet tested this DNA chromatography method with highly contaminated nucleotide samples extracted directly from sputum or skin specimens. Future improvements to further increase the speed of this method will focus on direct detection from tissue specimens without culturing.

In the present work, we sought to detect associations between indel regions and the subspecies, and between genetic loci and the *erm*(41) T28C mutation. Our method successfully discriminated the subspecies and inducible macrolide susceptibility of almost all of the clinical isolates, but the few exceptions represent a limitation of the method that should be considered. MabMT37 yielded a 1,200 bp and 450 bp product for locus MAB_2613 and MAB_1655 (similar to *M. bolletii* strains) and a 330 bp product for MAB_4665 (similar to *M. abscessus* strains). However, WGS-based phylogenetic and ANI analyses unambiguously categorized MabMT37 as *M. abscessus* (Table S1). We mapped raw sequence reads of MabMT37 to the *M. abscessus* ATCC 19977 genome and confirmed that there was no heterogeneity among the MAB2613, MAB_1655, and MAB_4665 loci (data not shown), suggesting that MabMT37 was likely a mono-clonal isolate. This result also suggested that, with respect to these genetic markers, MabMT37 had a chimeric genome structure between *M. abscessus* and *M. bolletii*. MabMT19 showed inducible clarithromycin resistance and had no mutation at *erm*(41) position 28, but it did cluster with other *erm*(41) T28C mutants and was positive for the primer set MAB18036_2558F and MAB18036_2558R (Table S1), suggesting that MabMT19 had a chimeric genome between *M. abscessus* clades. These observations are supported by a previous genomic study describing an asymmetrical gene flow between MABC subspecies that resulted in a highly mosaic genome architecture(50). Since our assays do not directly analyze housekeeping genes or the *erm*(41) gene itself, our approach could be affected by genome architecture mosaicism arising via horizontal gene transfer. To address this limitation, accumulation of genomic information for MABC clinical isolates from across the world and successive adjustments of target genomic sequences will be important.

In conclusion, we developed a rapid, easy-to-use, and accurate assay to identify subspecies and macrolide susceptibility of MABC by analyzing WGS data of clinical isolates. This assay could be introduced into clinical laboratory practice to facilitate selection of effective treatments, development of assays having improved discrimination and diagnostic capacity, and acquisition of precise, nation-wide epidemiological information for MABC. Although the incidence of the macrolide-susceptible *erm*(41) T28C mutation is unknown, a population of *M. abscessus* in our Japanese sample set (18.5%) carried this mutation. Phylogenetic relationships between these mutants and global circulating clones of *M. abscessus*(30) should be addressed in future studies. Additional international corroboration studies based on epidemiological and population genomic approaches will be required to address these fundamental research questions.

## Acknowledgments

The clinical isolates used in this study were sent from the hospitals and universities listed below. We appreciate the work of all of the clinicians at the following institutions who cared for patients infected with these mycobacteria: Hokkaido Social Insurance Hospital, Fukujuji Hospital Japan Anti-Tuberculosis Association (JATA), Saitama Medical University Hospital, National Hospital Organization (NHO) Tokyo Hospital, Showa University Fujigaoka Hospital, Keio University Hospital, National Defense Medical College Hospital, Kyorin University Hospital, Nagaya City University Hospital, NHO Minami-Kyoto Hospital, Kyoto Prefectural University of Medicine Hospital, Osaka City Northern-City Hospital, Osaka Hospital JATA, NHO Kinki-Chuo Chest Medical Center, NHO Matsue Medical Center, Kawasaki Medical School, NHO Higashi-Hiroshima Medical Center, Kyosai-Yoshijima Hospital, Kyushu University Hospital, and NHO Omuta Hospital. Computations were performed in part on the NIG supercomputer at the ROIS National Institute of Genetics. We thank Ms. Maki Okuda, Sayaka Kashiwagi, and Ginko Kaneda for their assistance.

## Availability of data and materials

All raw data are available by request to the corresponding authors.

## Author contributions

Strain collection: JYC, PRH, KM, AK, NH, SMitara Study conception: YH

Experimental design MY, YH

Performed experiments MY, SS, HF, MS, YM, SMiyam

Data analysis: MY, YH

Data interpretation: MY, YH

Figure production: MY Manuscript writing: MY, YH

Manuscript editing: MY, SS, JYC, HF, PRH, TA, KM, NH, SMitara, MA, YH

## Online Data Supplement

### Supplemental Methods

#### DNA sequencing and genomic analysis

Genomic DNA was extracted from each isolate with a NucleoSpin Plant II kit (MACHEREY-NAGEL, Düren, Germany) in accordance with the manufacturer’s instructions, and was used for Nextera XT library construction and genome sequencing with Illumina NovaSeq or MiniSeq. All raw read data for the newly sequenced strains (148 strains isolated in Japan) in this study were deposited in the DNA Data Bank of Japan (DDBJ) and mirrored at the National Center for Biotechnology Information (NCBI) under BioProject accession number PRJDB10333.

The Illumina read data for each isolate were *de novo* assembled into contigs using a Shovill pipeline with default settings (https://github.com/tseemann/shovill). Table S3 lists the number of contigs, raw coverage and N50 value of each isolate. We combined the data with the complete genome sequence of *M. abscessus* ATCC19977(1), *M. massiliense* JCM 15300(2), and *M. bolletii* BD(3). Phylogenetic analysis using a core gene set from the MABC isolates from Japan was next performed. We annotated each genome using DFAST-core ver.

1.0.3 with the default setting(4) and the resulting gene annotations (GFF3 format) were used with Roary software and the “-cd 100” option(5) to compute core-gene alignments for the isolates. Using the 2,827,548 bp-long core-gene alignment carrying 62,196 SNP positions, we estimated a maximum-likelihood tree using RAxML ver. 8.2.12(6). We used the following parameters that indicate the GTR + G4 model of DNA substitution with estimation of the shape parameter of the gamma distribution by maximizing the likelihood: -f a -m GTRGAMMA. Average nucleotide identity (ANI) values among all MABC isolates were calculated using fastANI with default settings(7). To identify single nucleotide polymorphisms (SNPs) in MABC isolates, we used MUMmer to conduct pairwise genome alignment between *M. abscessus* ATCC 19977 and one of the 150 strains including *M. bolletii* BD and *M. massiliense* JCM 15300(8). We then used an in-house Perl script to combine the alignments in a multiple whole-genome alignment, in which each position corresponded to that of the ATCC 19977 genome. The multiple whole-genome alignments were used for extracting and investigating the *erm*(41) genotype (MAB_2297, nucleotide position: 2345955 to 2346476 in ATCC 19977^T^) of all strains using SeqKit(9).

#### Construction of primers for discriminating MABC subspecies and the *erm*(41) T28C sequevar

To construct primer sets for discriminating MABC, we first aligned the complete genome sequences of *M. massiliense* JCM15300 and *M. bolletii* BD to the *M. abscessus* ATCC19977 reference genome and draft genomes of representative 14 clinical isolates using progressiveMauve and then visually identified unique insertion/deletion (indel) regions in each of the three subspecies. Three primer sets were designed around the indel regions to allow differentiation of the subspecies based on PCR amplicon size (Table 1). To construct primer sets to determine the sub-clades of the *M. abscessus erm*(41) T28C sequevar, we identified accessory genes that were significantly associated with the lineage using Scoary(10). Primer sets were designed around the accessory genes used as genetic markers of the lineage (Table 2).

### Statistical analysis

Statistical analyses were performed with R software (www.r-project.org). The R function binom.test() was used to assess 95% confidence intervals (95% CIs) of agreement rates between the multiplex PCR/DNA chromatography and WGS-based identification. Differences in accuracy between the multiplex PCR developed in this study and our previous multiplex PCR were statistically assessed using the R function prop.test().

### Ethics Statement

This study was reviewed and approved by the medical research ethics committee of the National Institute of Infectious Diseases for inclusion of human subjects (#1004 and #1005 for the Japanese and Taiwanese, respectively).

## Supplemental Figure Legends

**Figure S1.**
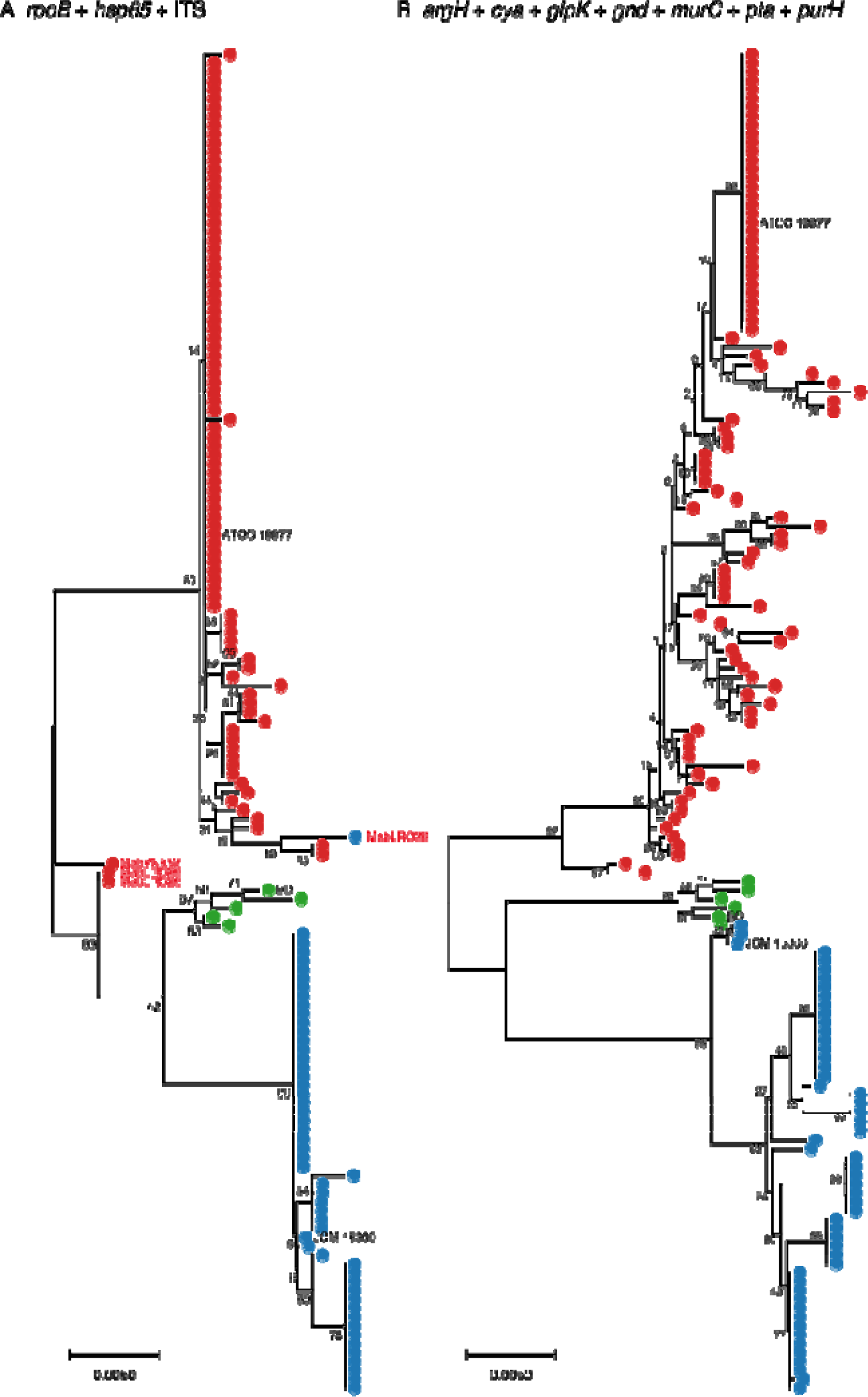
Maximum-likelihood trees constructed from concatenated sequences. **A.** Concatenated sequences of *rpoB* (409-bp), *hsp65* (409-bp), and ITS (298-bp) fragments to differentiate subspecies of MABC as previously described (11). Samples having results that differed from those for WGS-based analyses are highlighted. **B.** Concatenated sequences of 7 housekeeping genes [*argH* (480 bp), *cya* (510 bp), *glpK* (534 bp), *gnd* (480 bp), *murC* (537 bp), *pgm* (495 bp), *pta* (486 bp), and *purH* (549 bp)] to differentiate subspecies of MABC as described previously(12). Sequences were extracted from the whole-genome alignment of each isolate to the reference strain *M. abscessus* ATCC 19977 (see Supplemental Methods). The trees for all isolates (n=148) and three reference strains (*M. abscessus* ATCC 19977, *M. massiliense* JCM 15300, and *M. bolletii* BD) were computed using MEGA ver. 10.1.7 with 500 bootstrap replicates. The bootstrap support values (%) are indicated for each node. Colored circles correspond to each member of MABC determined by WGS-based analyses (**Fig. 1** and **Fig. S2**): red, *M. abscessus*; blue, *M. massiliense*; green, *M. bolletii*, respectively.

**Figure S2.**
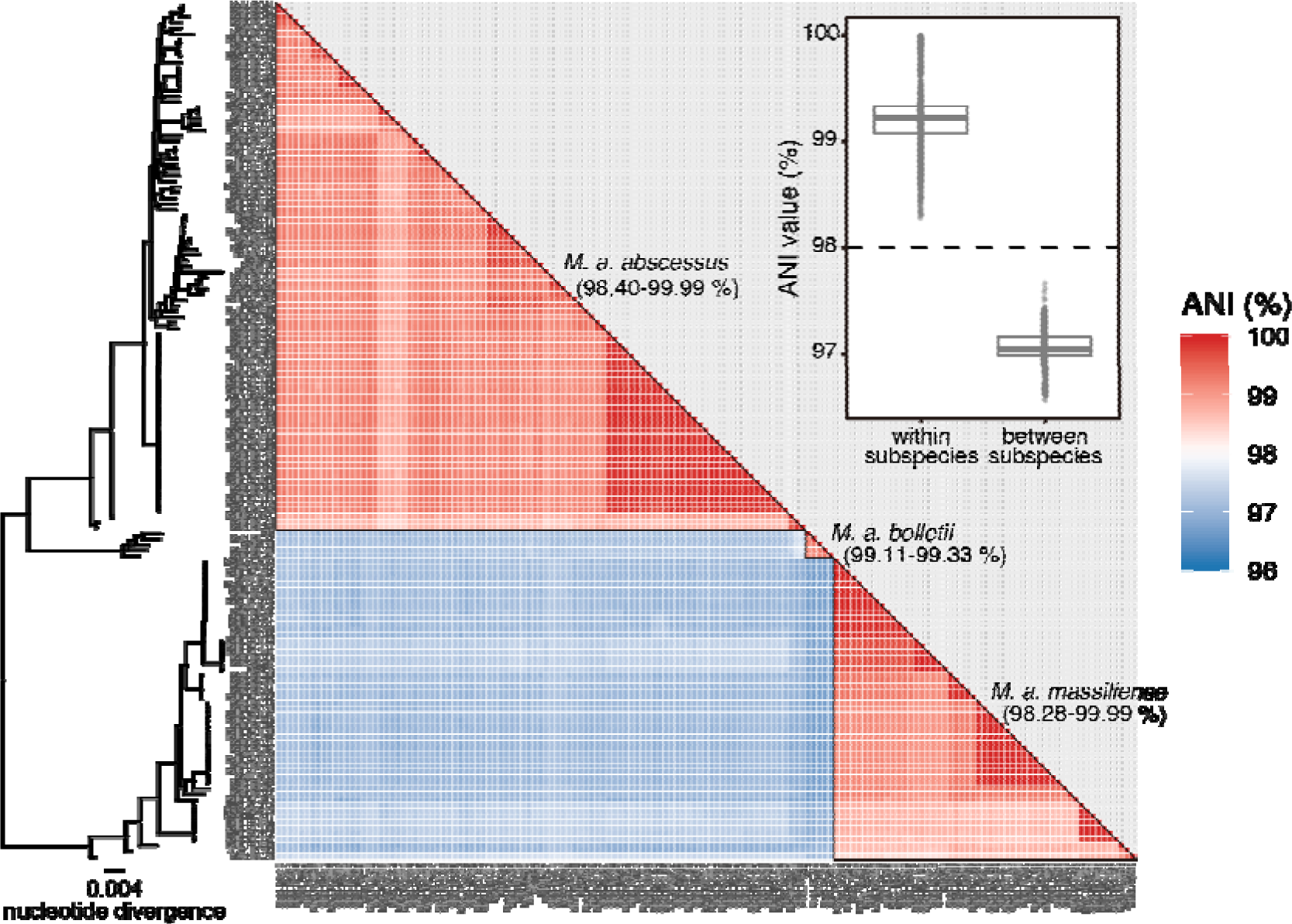
Average nucleotide identity matrix of MABC isolates. Average nucleotide identities among 148 isolates and three reference strains (*M. abscessus* ATCC 19977, *M. massiliense* JCM 15300, and *M. bolletii* BD) were measured for all strain pairs using fast ANI(7). Red and blue boxes indicate ANI values higher or lower than 98% for corresponding strain pairs. A boxplot indicates the distribution of ANI values within or between subspecies of MABC. The maximum likelihood tree corresponds to that depicted in **Fig. 1**.

**Figure S3.**
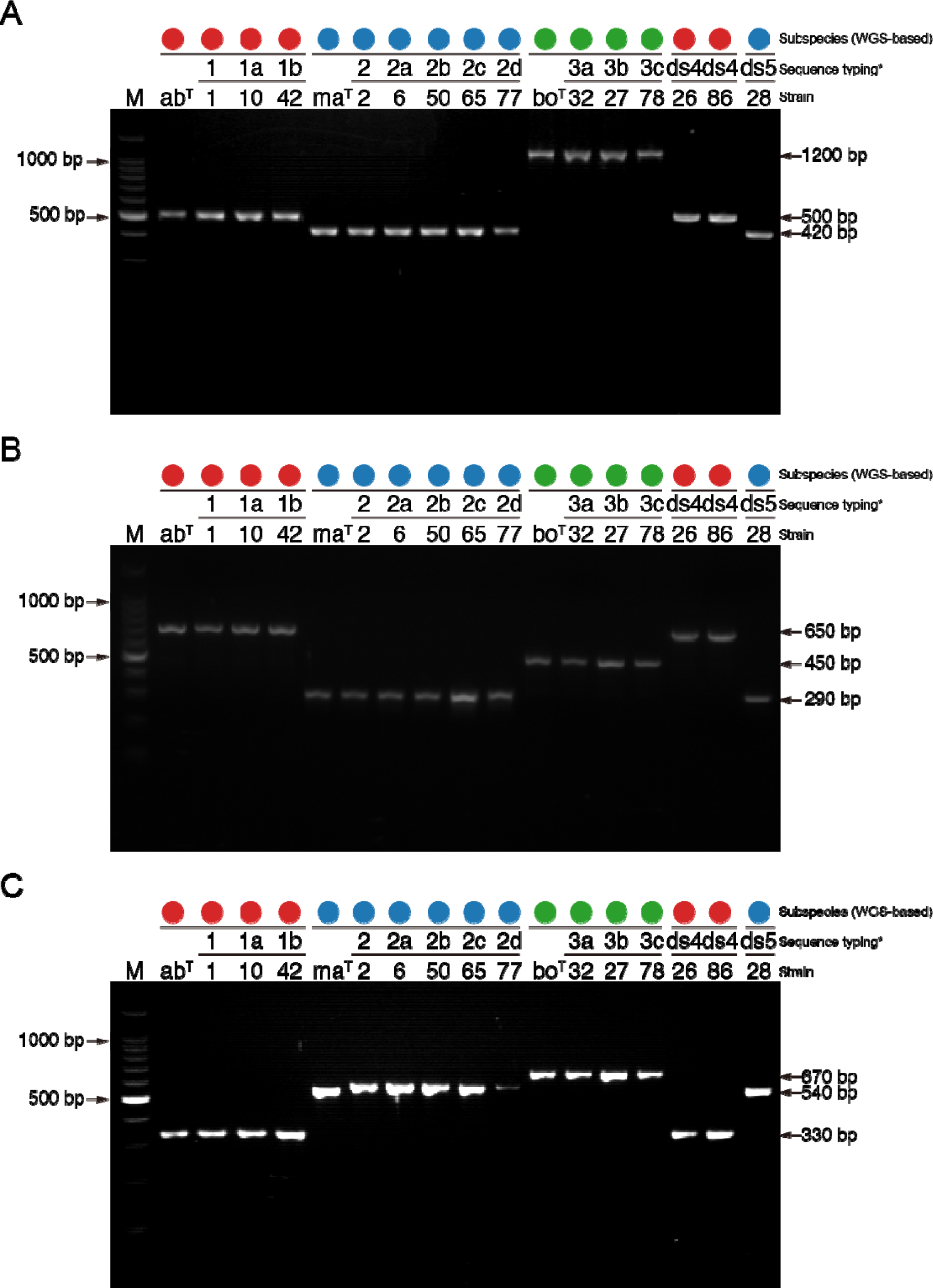
Representative single PCR to differentiate MABC subspecies. PCR was performed with primer pair **A.** MAB2613F and MAB2613R, **B.** MAB_1655F and MAB_1655R, and **C.** MAB_4665F and MAB_4665R. Types 1, 1a, 1b, 2, 2a, 2b, 2c, 2d, 3, 3a, 3b, 3c, 4 and 5 are the sub-groupings (sequevars) of the clinical isolates based on their sequences [Table 3 of the previous article]. Numbers below the sequevars correspond to reference strains (ab^T^, ma^T^, or bo^T^) or strain numbers of clinical isolates described in the previous article(11). Lanes: M, DNA marker (100 bp ladder); ab^T^, *M. abscessus* ATCC19977); ma^T^, *M. massiliense* (JCM15300); bo^T^, *M. bolletii* (JCM15297). Colored circles correspond to each member of MABC determined by WGS-based analysis (**Fig. 1** and **Fig. S2**): red, *M. abscessus*; blue, *M. massiliense*; green, *M. bolletii*, respectively. The PCR reaction products were electrophoresed on 2% agarose gels.

**Figure S4.**
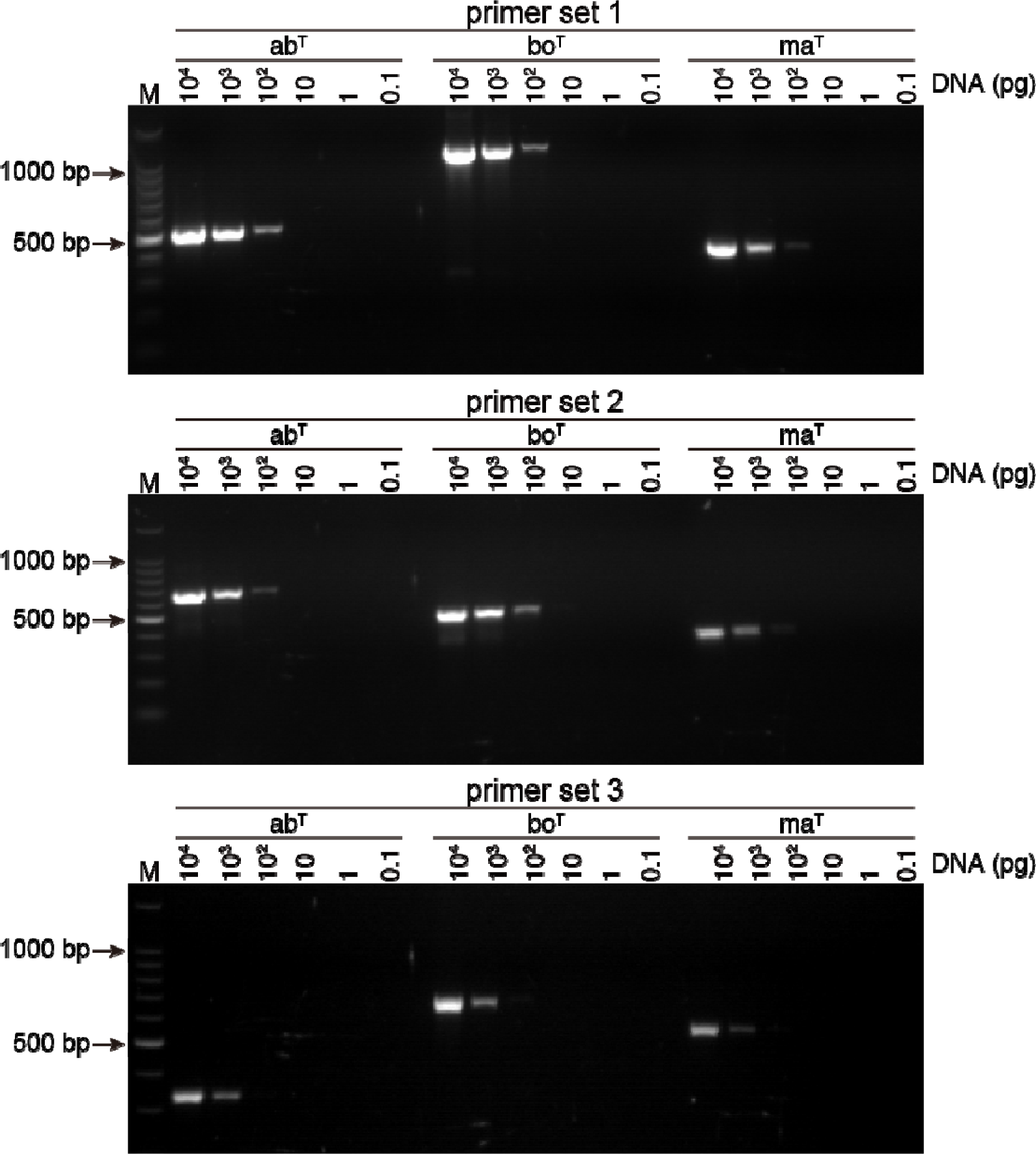
Sensitivity test of PCR primers to differentiate MABC subspecies. The sensitivity of primers to differentiate MABC subspecies was tested using 10 μg undiluted or ten-fold diluted DNA from the three MABC reference strains. Primer set 1, primer pair MAB2613F and MAB2613R; Primer set 2, primer pair MAB_1655F and MAB_1655R. Lanes: M, DNA marker (100 bp ladder); ab^T^, *M. abscessus* ATCC19977); ma^T^, *M. massiliense* (JCM15300); bo^T^, *M. bolletii* (JCM15297). The PCR reaction products were electrophoresed on 2% agarose gels.

**Figure S5.**
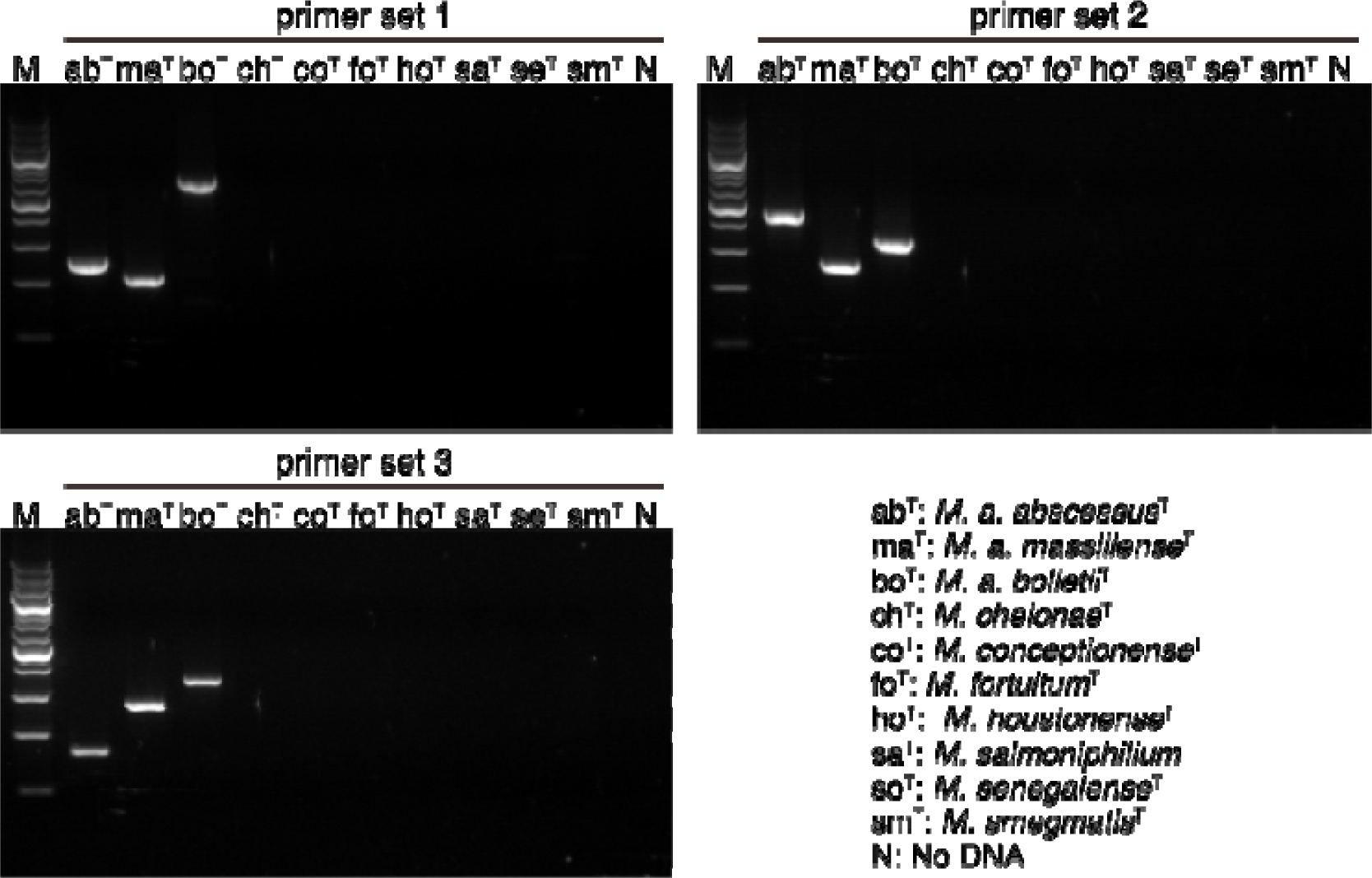
Specificity test of PCR primers to differentiate subspecies of MABC. The tests were performed with MAB2613F/MAB2613R (Primer set 1), MAB2613F/MAB2613R (Primer set 2), and MAB_1655F/MAB_1655R (Primer set 3), with 4 ng of each indicated mycobacterial genomic DNA. The PCR reaction products were electrophoresed on 2% agarose gels. M, DNA marker (100 bp ladder). All isolates other than the *M. abscessus* complex were negative in this assay [Only those representative data for rapid-growing NTM (ch^T^, *M. chelonae*^T^; co^T^, *M. conceptionense*^T^; fo^T^; *M. fortuitum*^T^; ho^T^, *M. houstonense*^T^; sa^T^, *M. salmoniphilium*^T^; se^T^, *M. senegalense*^T^; sm^T^, *M. smegmatis*^T^) are shown in this figure .

**Figure S6.**
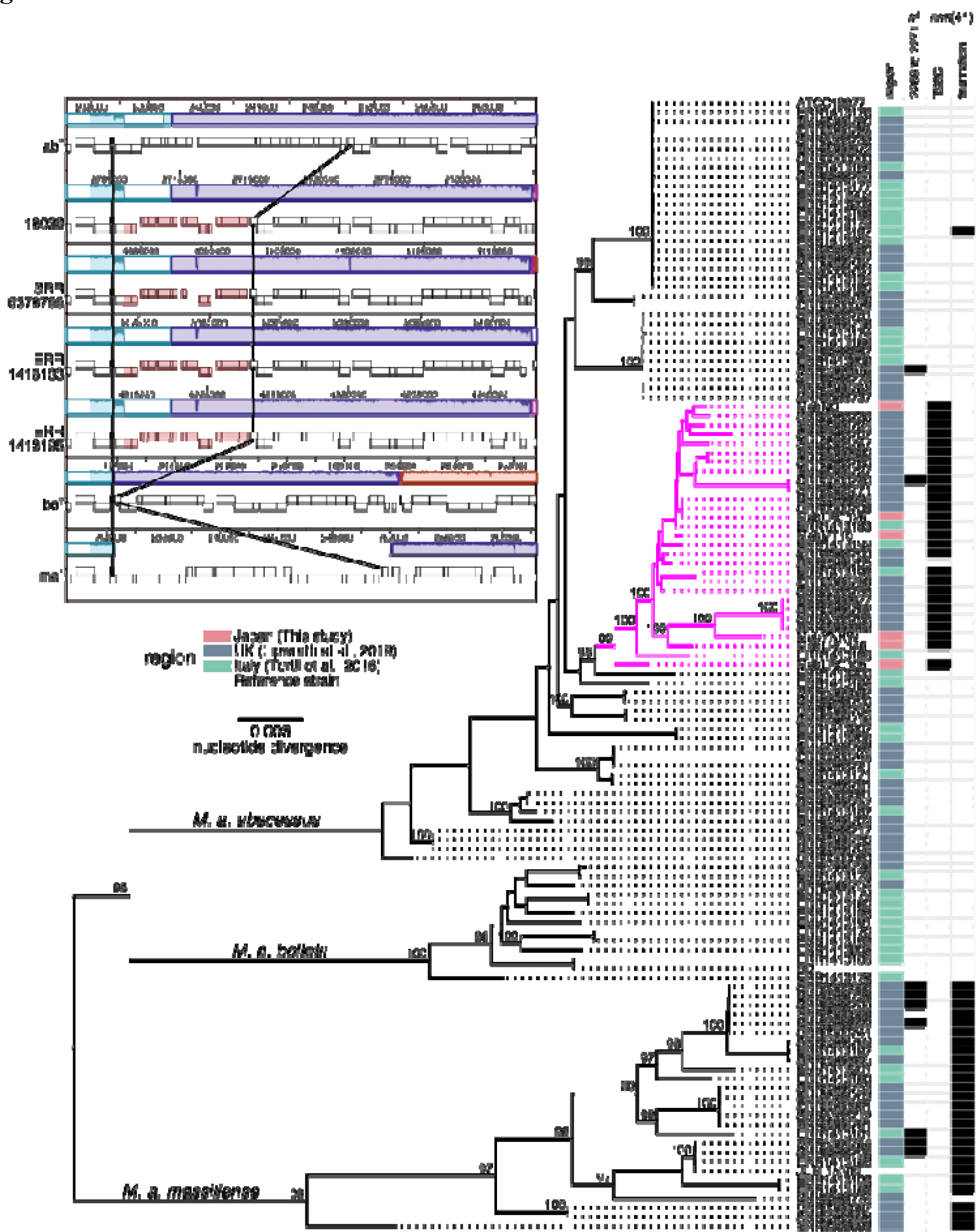
Maximum likelihood core-gene phylogeny of MABC from Japan, UK and Italy. Core-genome alignment of 120 isolates and three reference strains (*M. abscessus* ATCC19977, *M. massiliense* JCM 15300, and *M. bolletii* BD) of MABC were computed using Roary(5). The alignment containing 2,508 genes and 45,226 variable positions was used to construct a maximum likelihood tree using RAxML(6) with 300 bootstrap replicates. Bootstrap values > 90% for the major nodes are shown. The scale bar represents the mean number of nucleotide substitutions per site (SNP/site) on the respective branch. Samples are highlighted based on inclusion in three major clusters corresponding to subspecies of MABC. Colored boxes indicate countries where the corresponding strain was isolated: red, Japan; blue, UK; green, Italy; gray, reference strain. This tree is also annotated for the presence (black) and absence (light gray) of macrolide resistance-associated mutations. These mutations include the presence of a T-to-C substitution in position 28 or a truncation of the *erm*(41) gene that are associated with inducible resistance to macrolides; the presence of mutations at *rrl* position 2269 to 2271 confers acquired macrolide resistance. Substitutions or truncation with asterisks indicate non-synonymous mutations. The lineage to which all *M. abscessus erm*(41) T28C mutants belong is highlighted in magenta. **(inset) Visualization of lineage-specific genomic loci.** A progressiveMauve alignment of three clinical isolates carrying the *erm*(41) T28C mutation and the reference strains is shown. Red boxes indicate genomic regions specific to the lineage to which all *M. abscessus erm*(41) T28C mutants belong.

**Figure S7.**
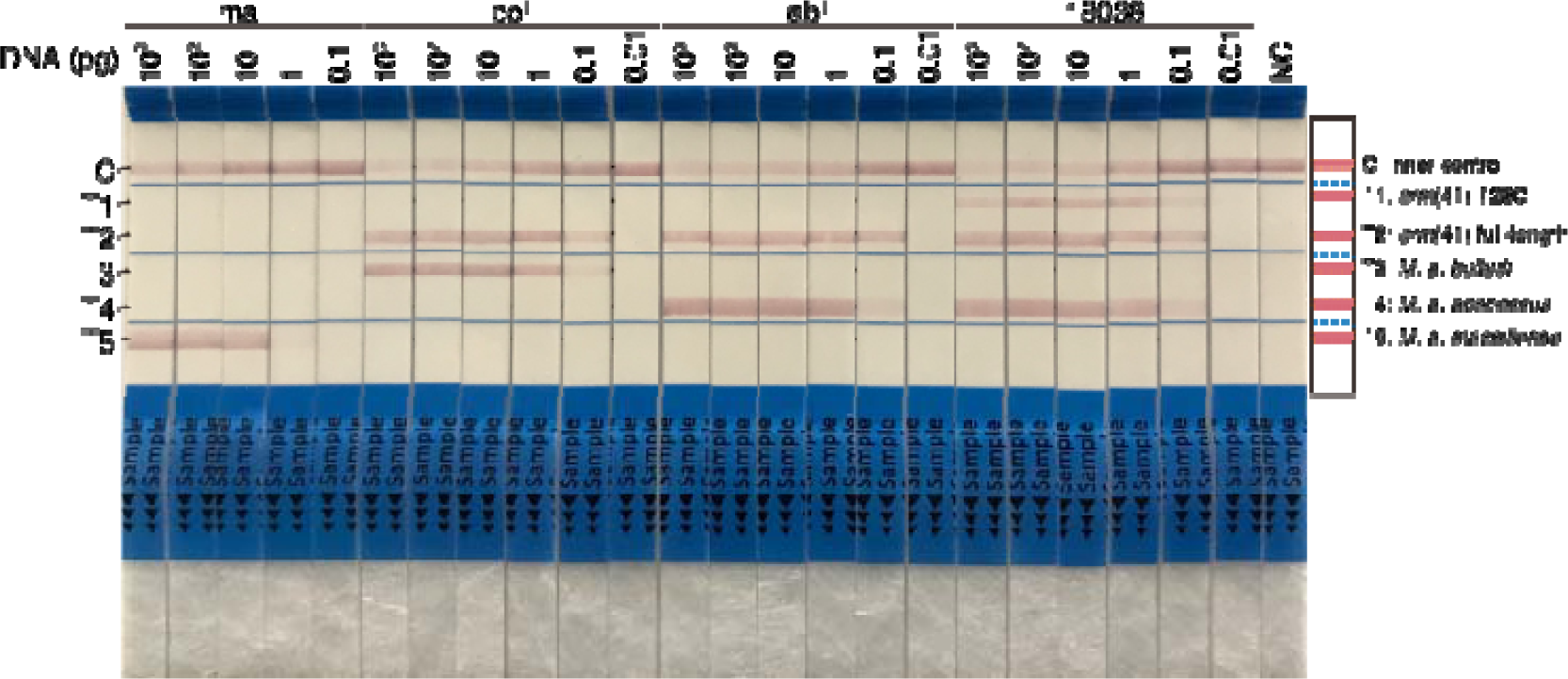
Sensitivity of the DNA chromatography method to differentiate subspecies and macrolide susceptibility resistance of MABC. A sensitivity test was performed with 1 ng undiluted or ten-fold diluted DNA from three reference strains and a clinical isolate carrying the *erm*(41) T28C polymorphism. Numbers correspond to type strains of MABC (ab^T^, ma^T^, or bo^T^) or the strain names of the clinical isolates (Mab18036). Bands: C, inner (negative) control; T1, *erm*(41) T28C polymorphism; T2, intact *erm*(41) genes; T3, *M. bolletii*; T4, *M. abscessus*; T5, *M. massiliense*.

**Figure S8.**
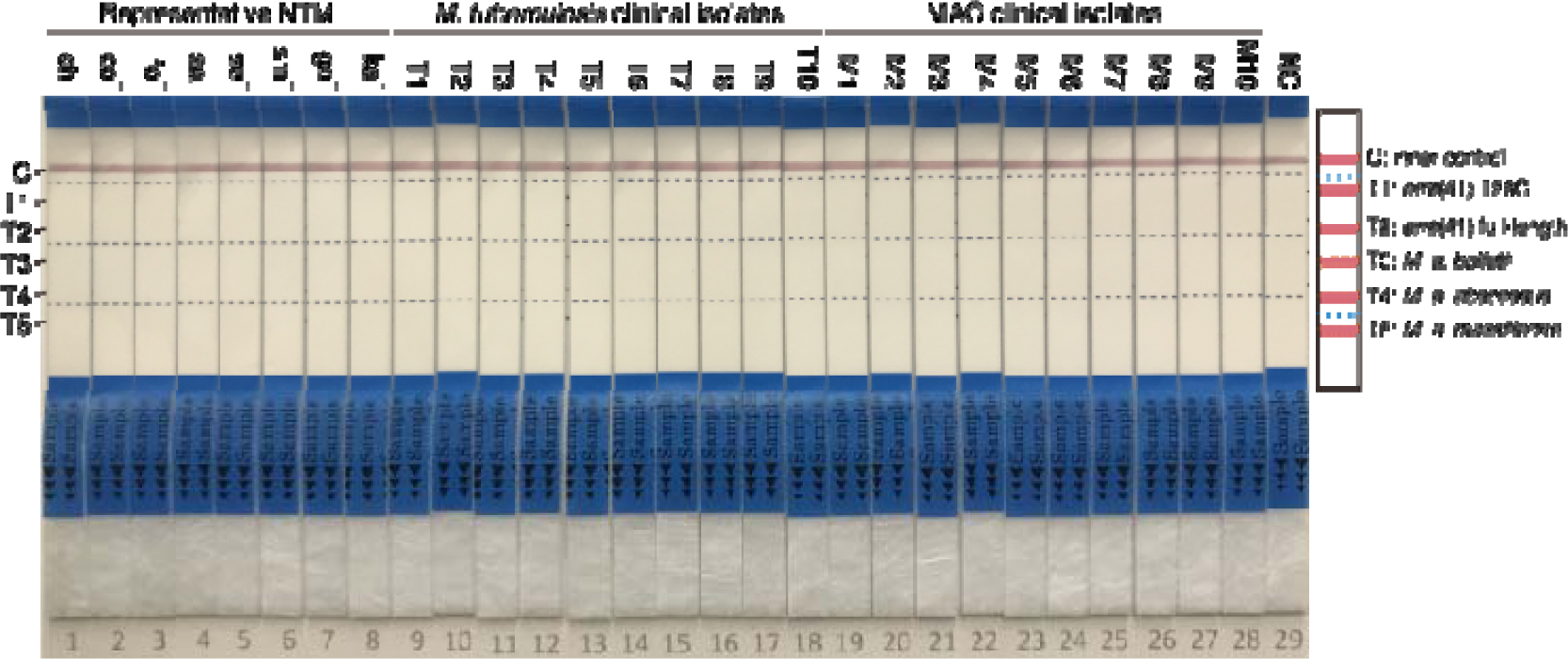
Specificity of the DNA chromatography method to differentiate subspecies and macrolide susceptibility of MABC. Specificity tests were performed using 4 ng of each indicated mycobacterial genomic DNA. Several laboratory and clinical isolates were tested in this assay. All isolates other than the *M. abscessus* complex were negative in this assay [Only representative data for NTM (ch^T^, *M. chelonae*^T^; co^T^, *M. conceptionense*^T^; fo^T^, *M. fortuitum*^T^, go^T^, *M. gordonae*^T^; ka^T^, *M. kansasii*^T^; sa^T^, *M. salmoniphilium*^T^; se^T^, *M. senegalense*^T^; sm^T^, *M. smegmatis*^T^) and *M. tuberculosis* and *M. avium* clinical isolates are shown in this figure].

## Supplementary Tables

**Table S1.**
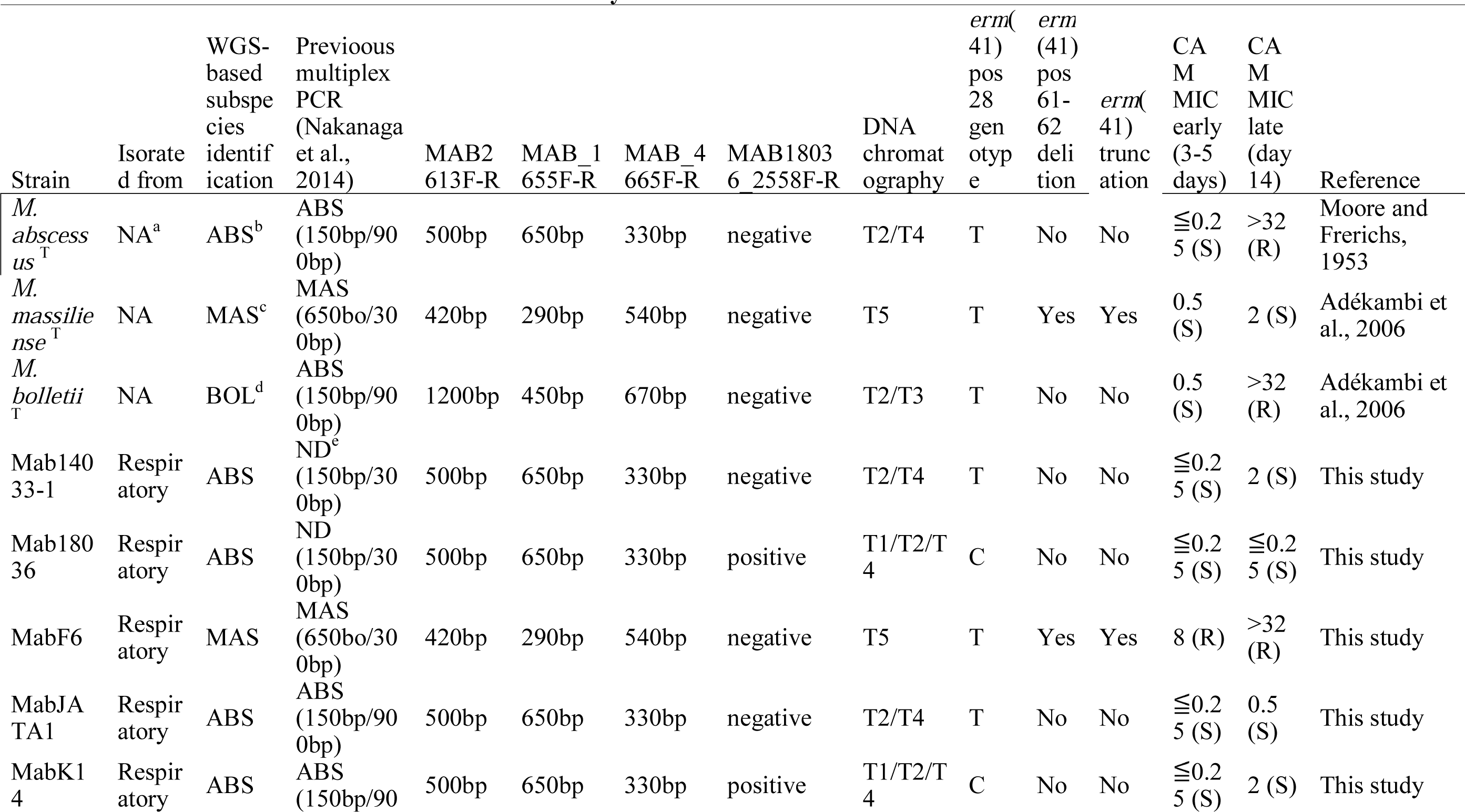
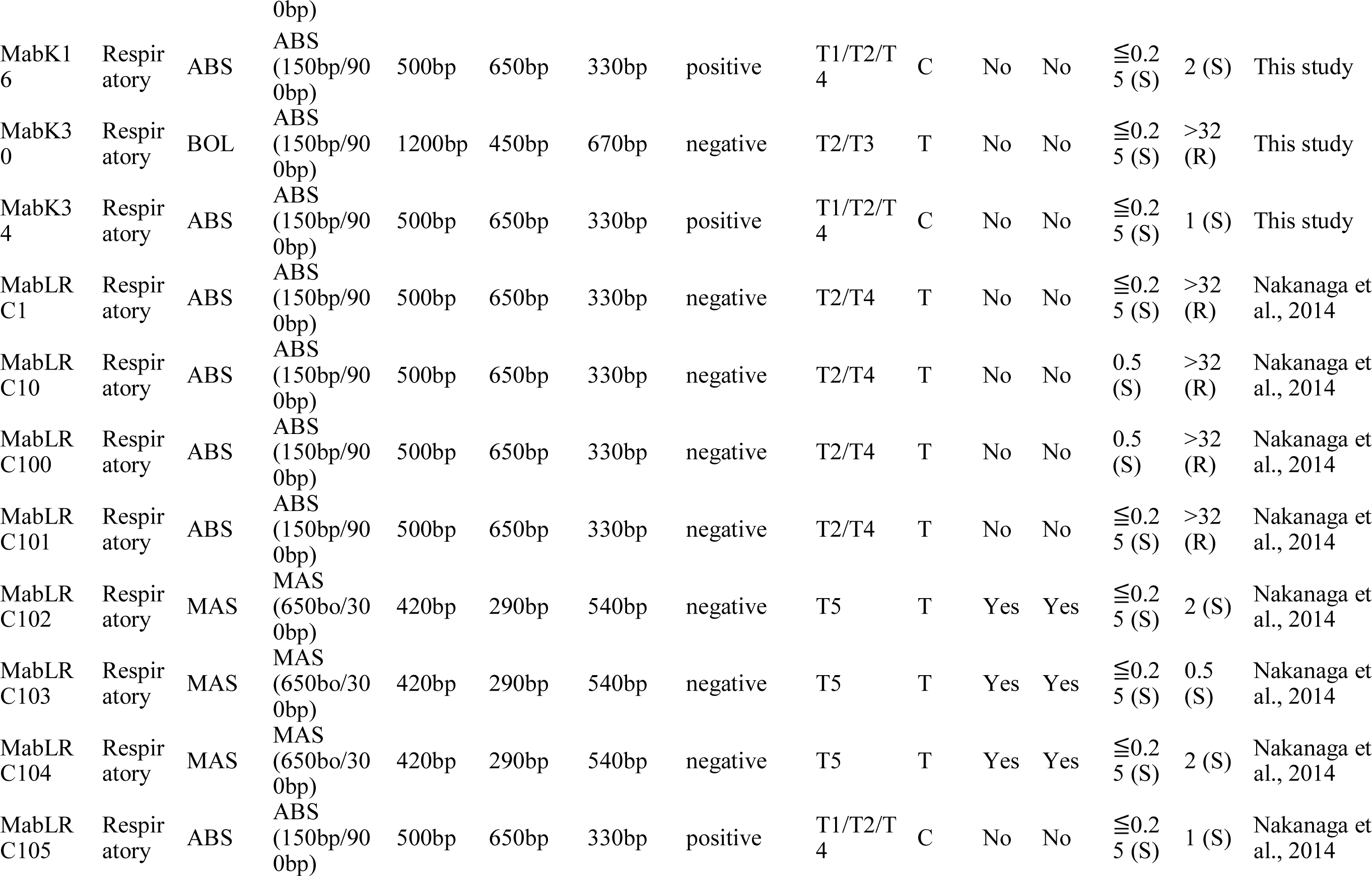
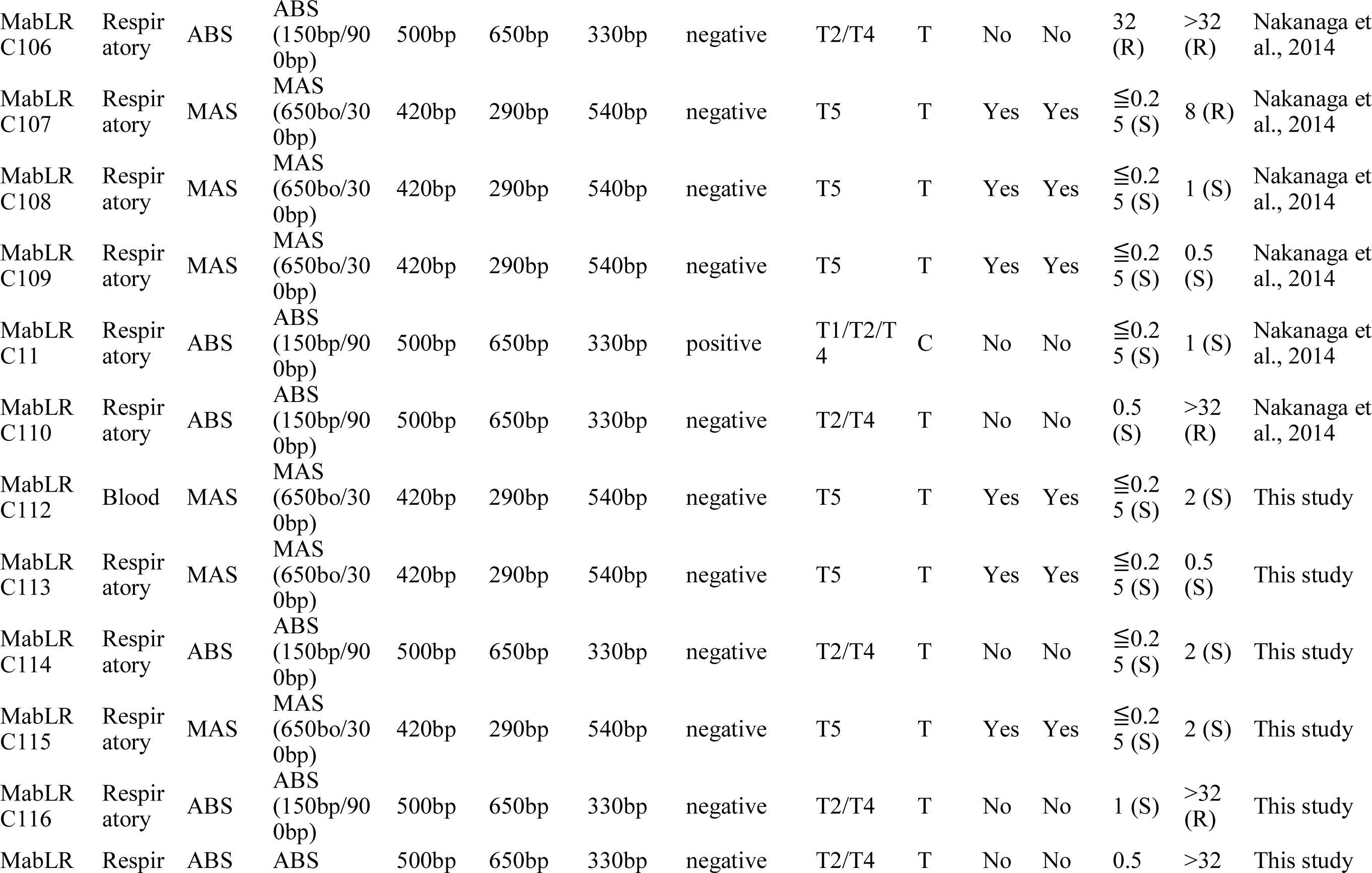
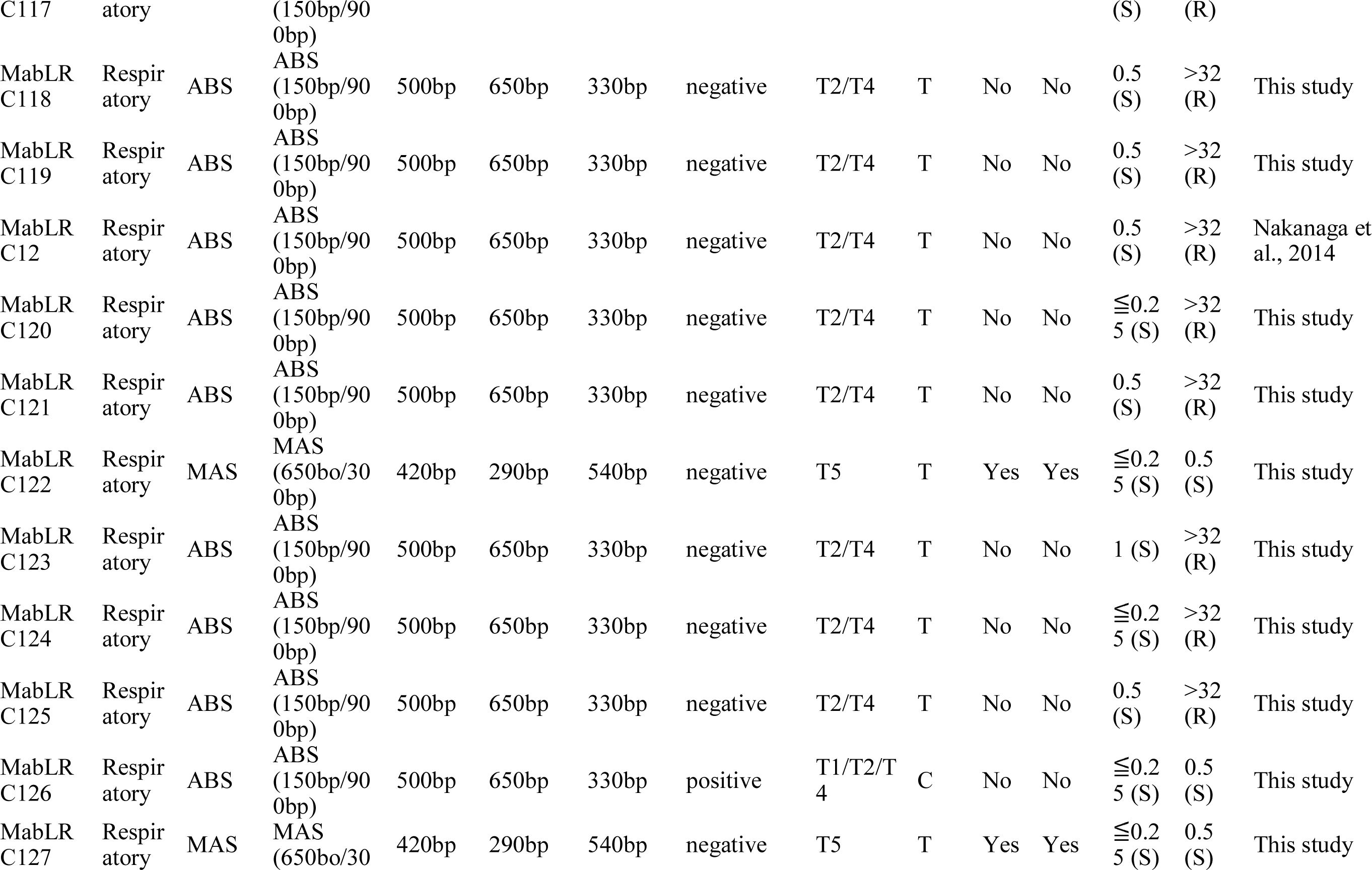
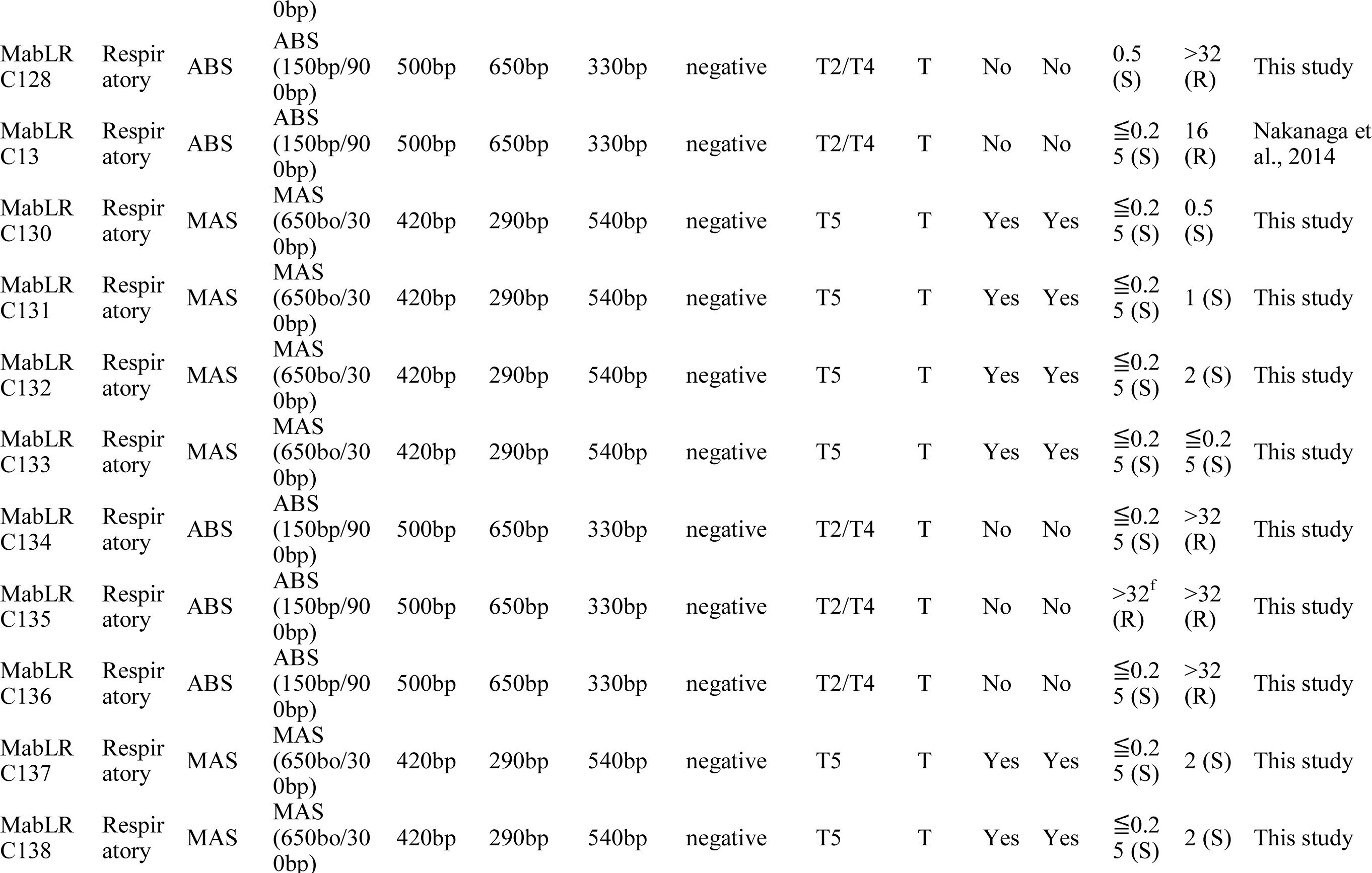
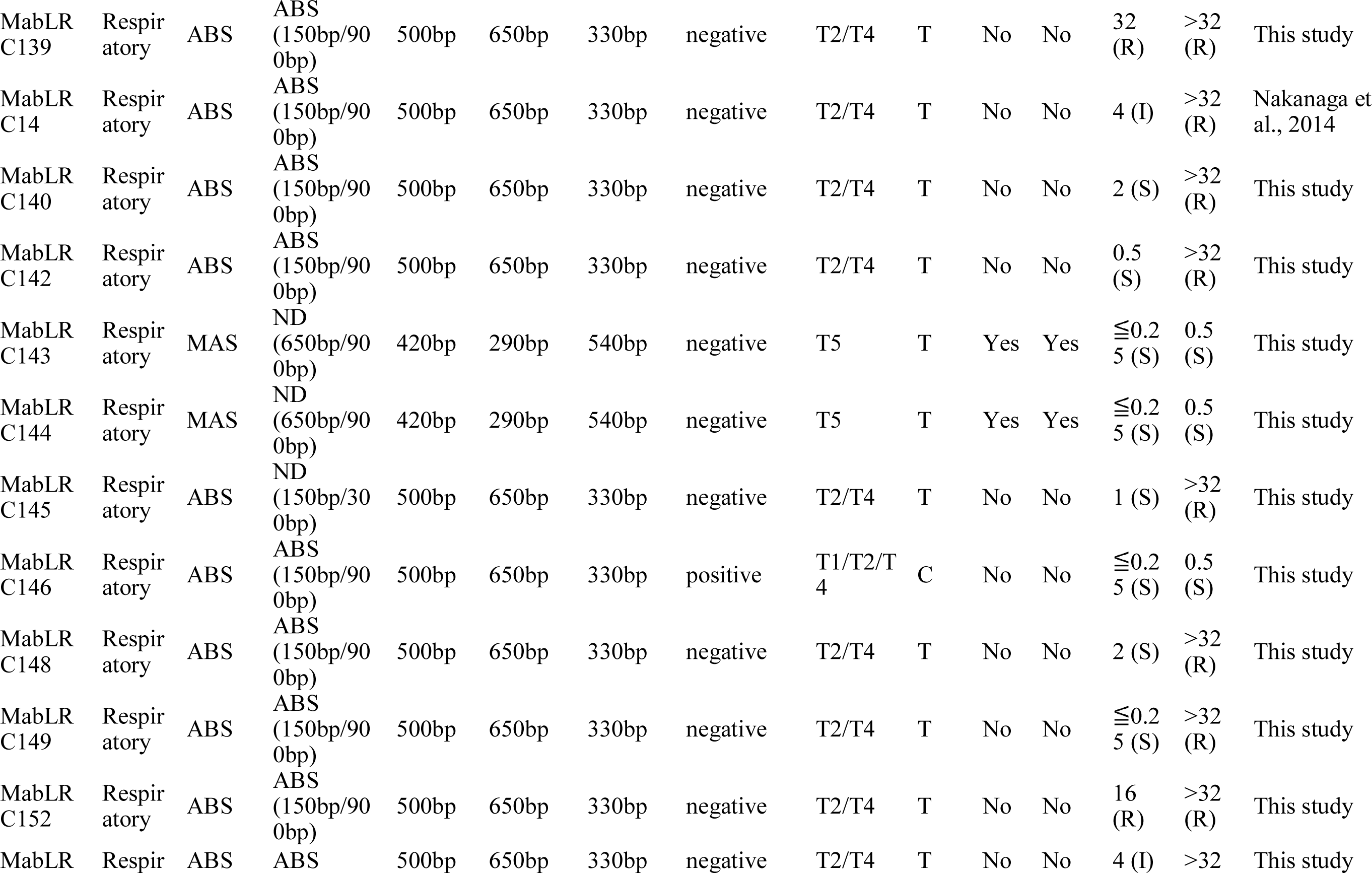
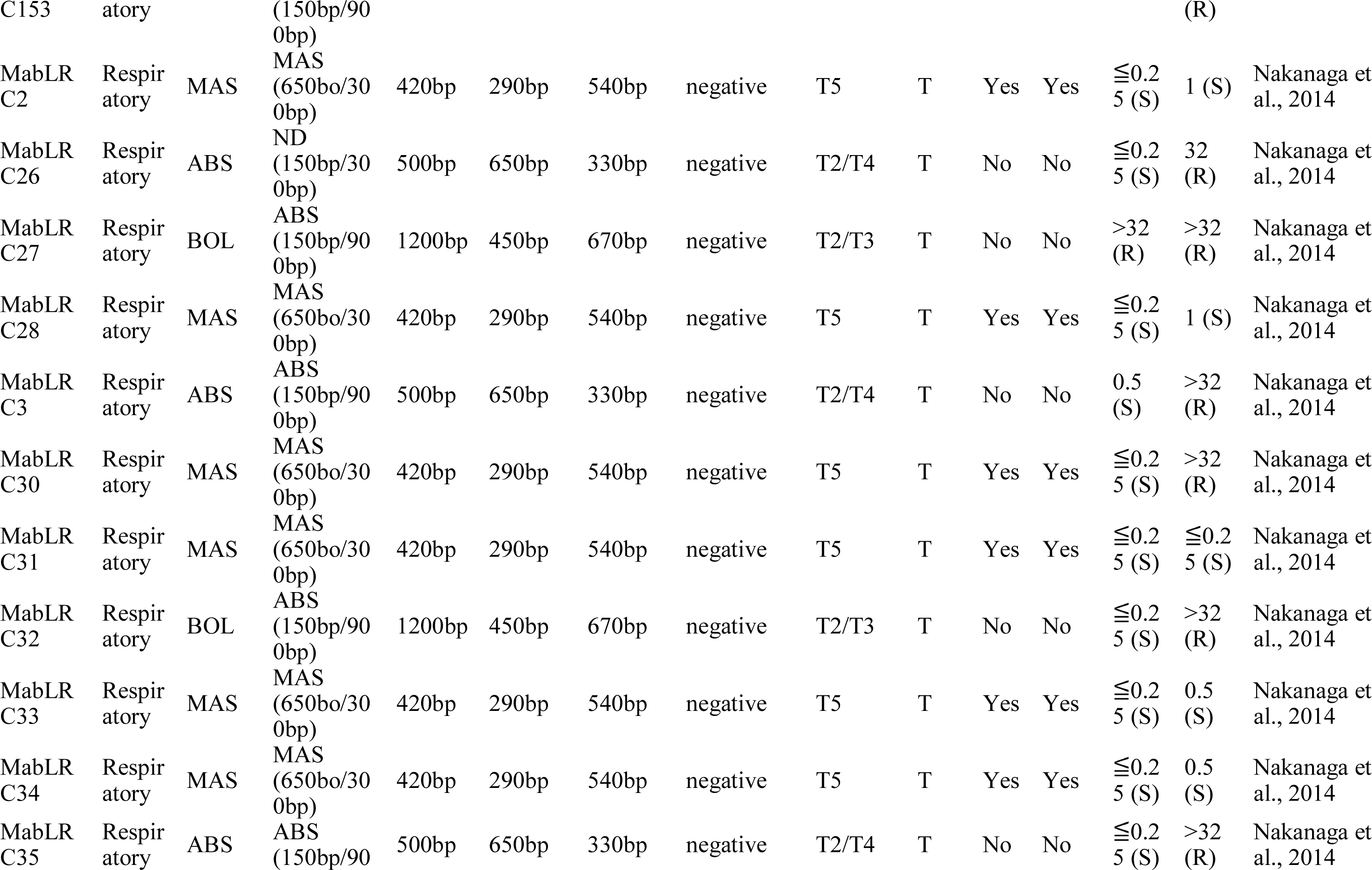
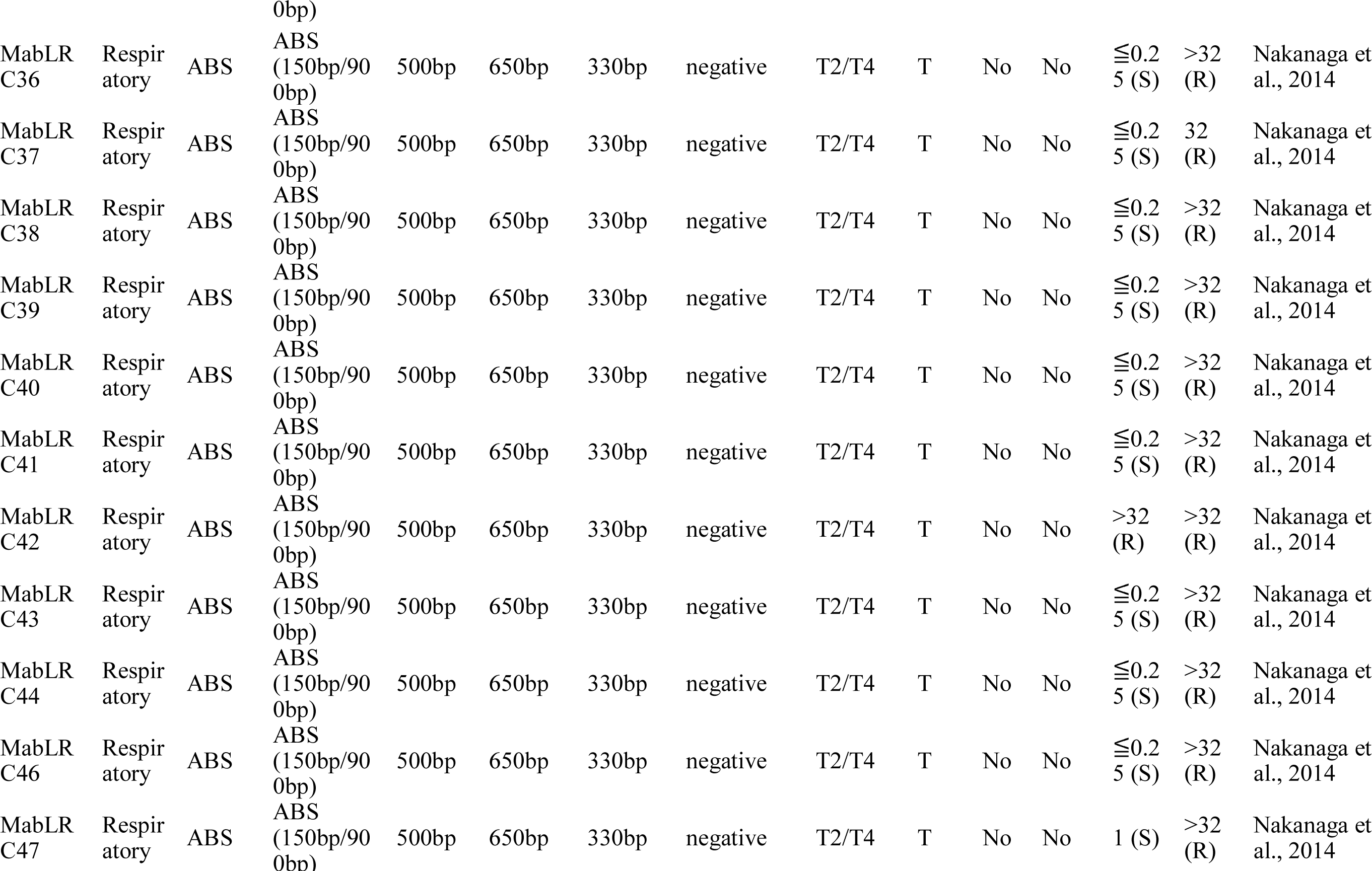
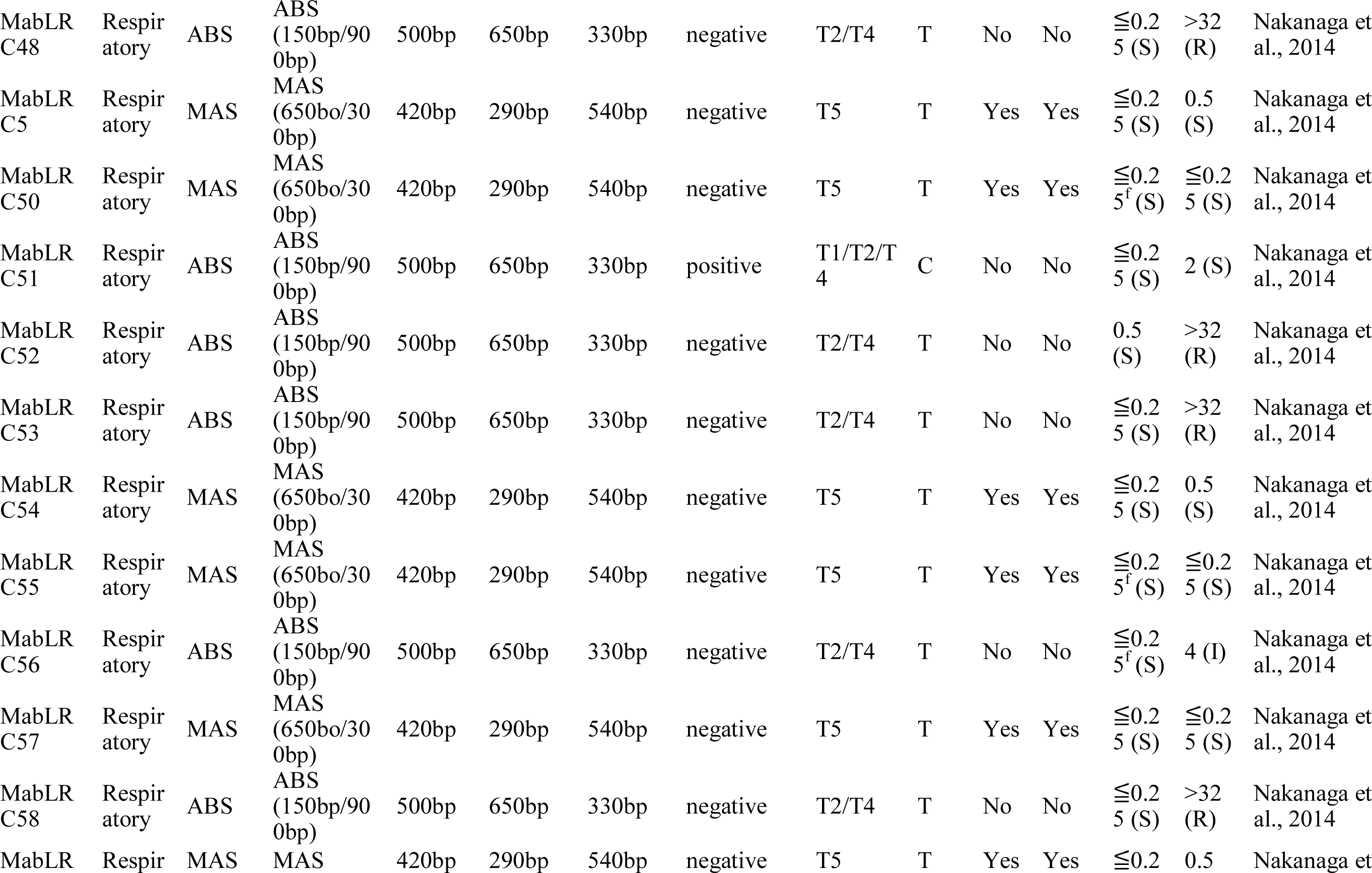
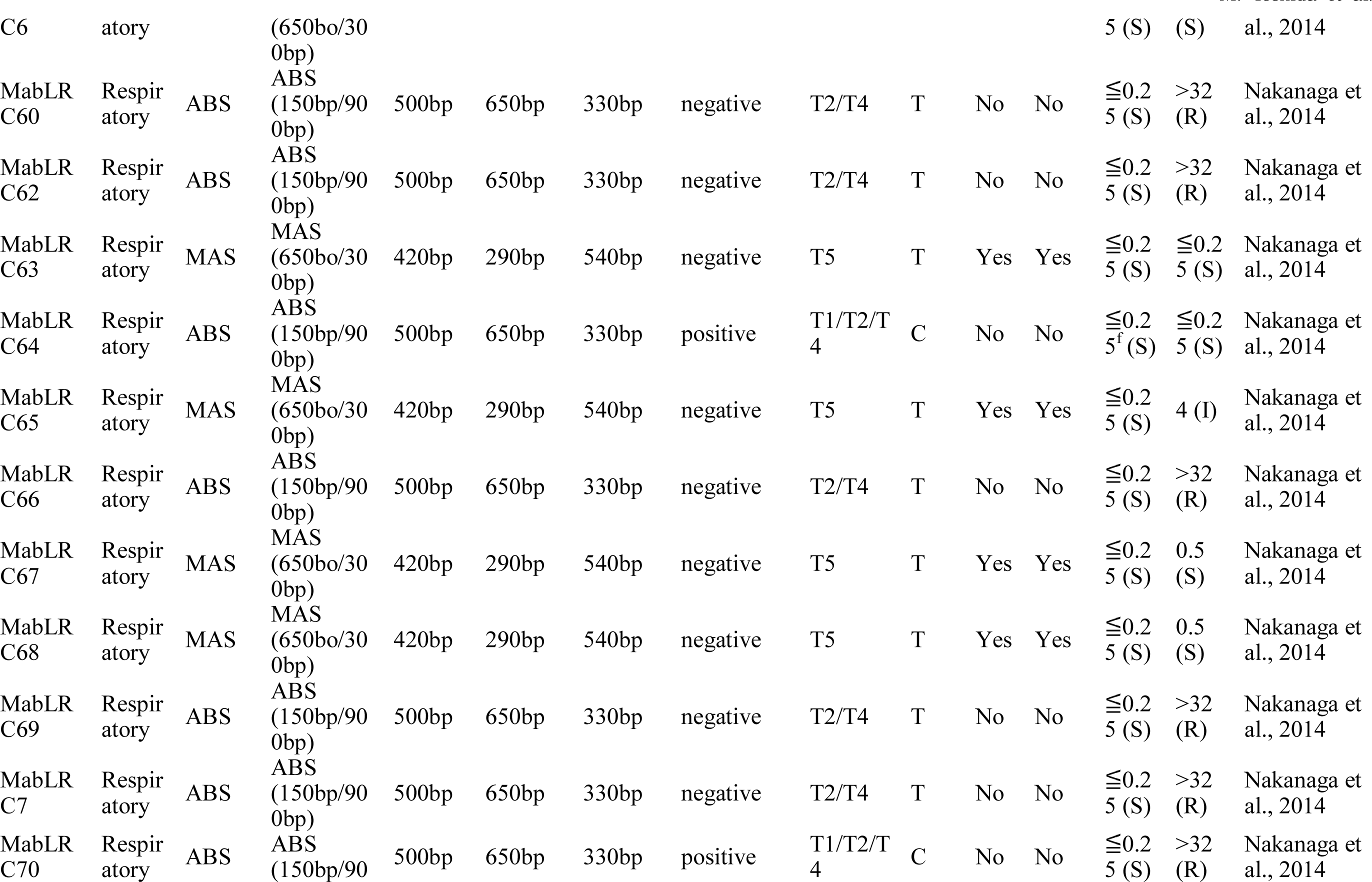
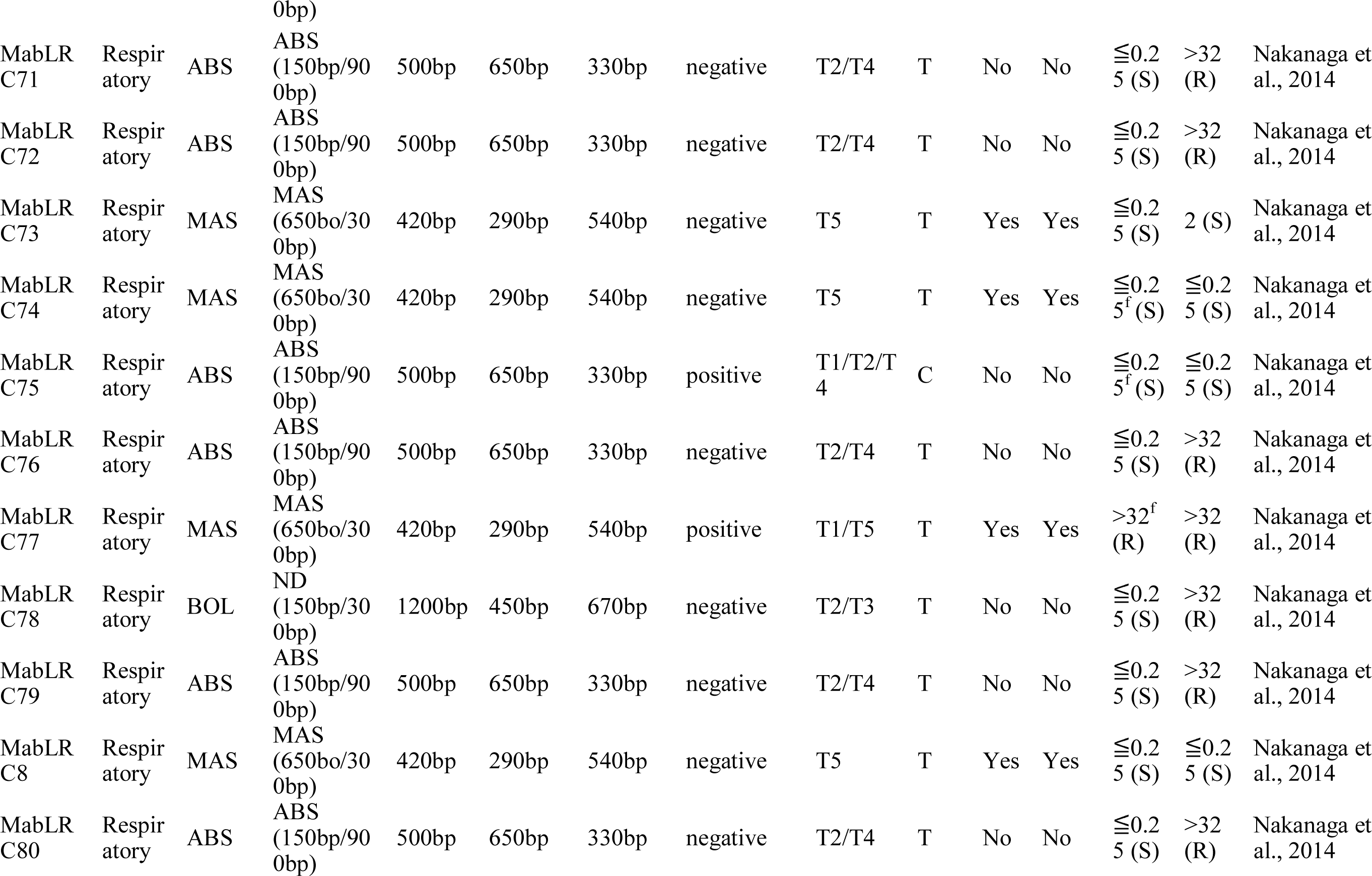
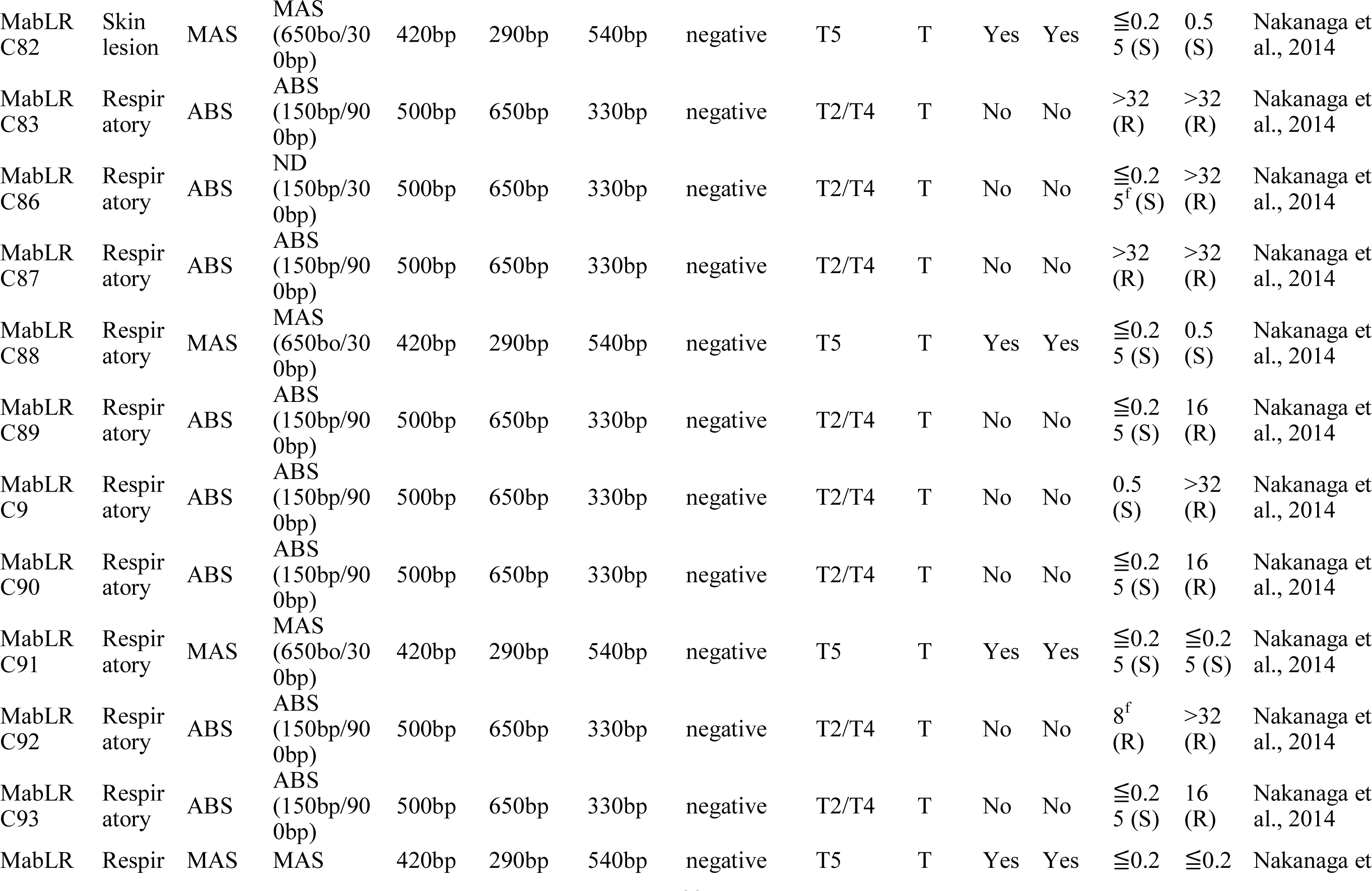
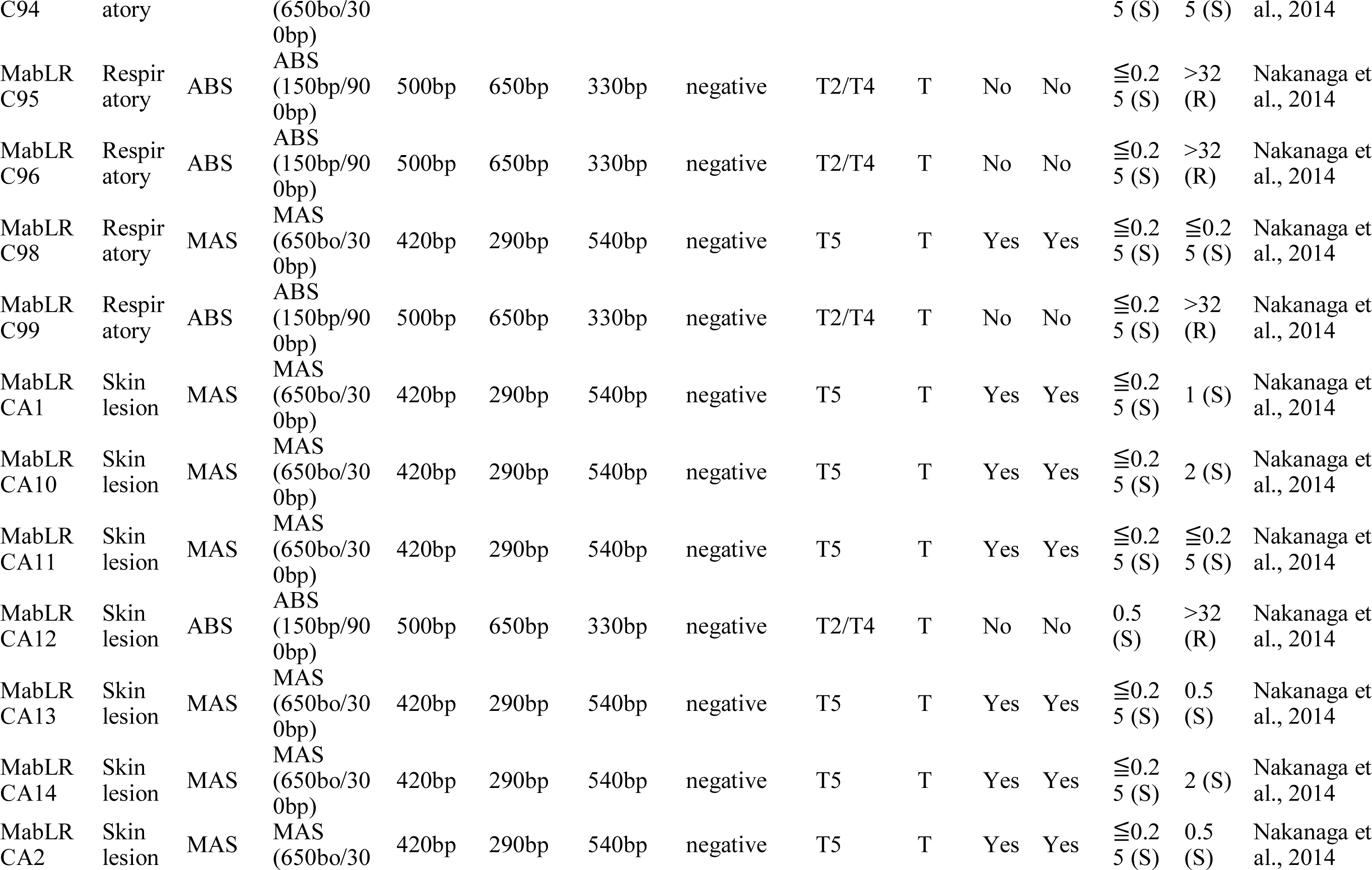
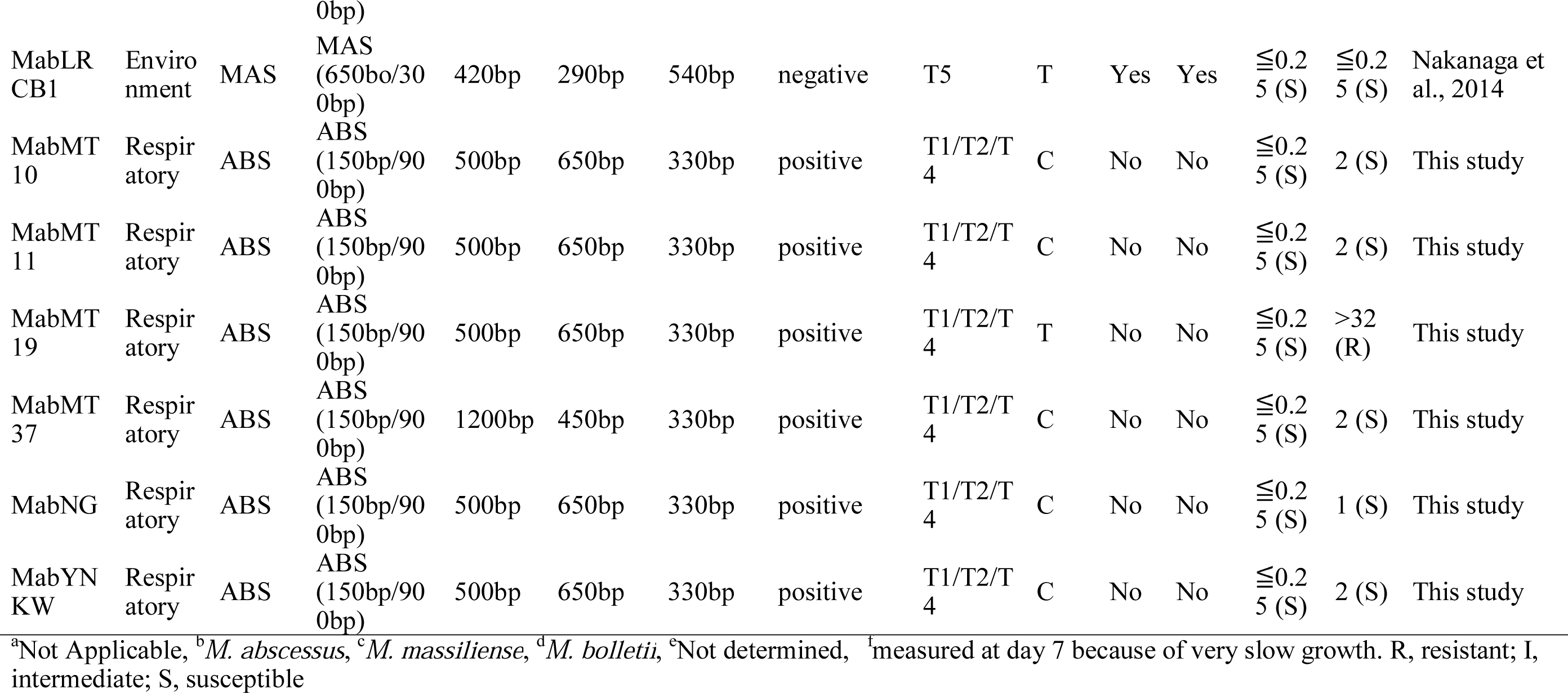
Bacterial strains and overall results of this study.

**Table S2.**
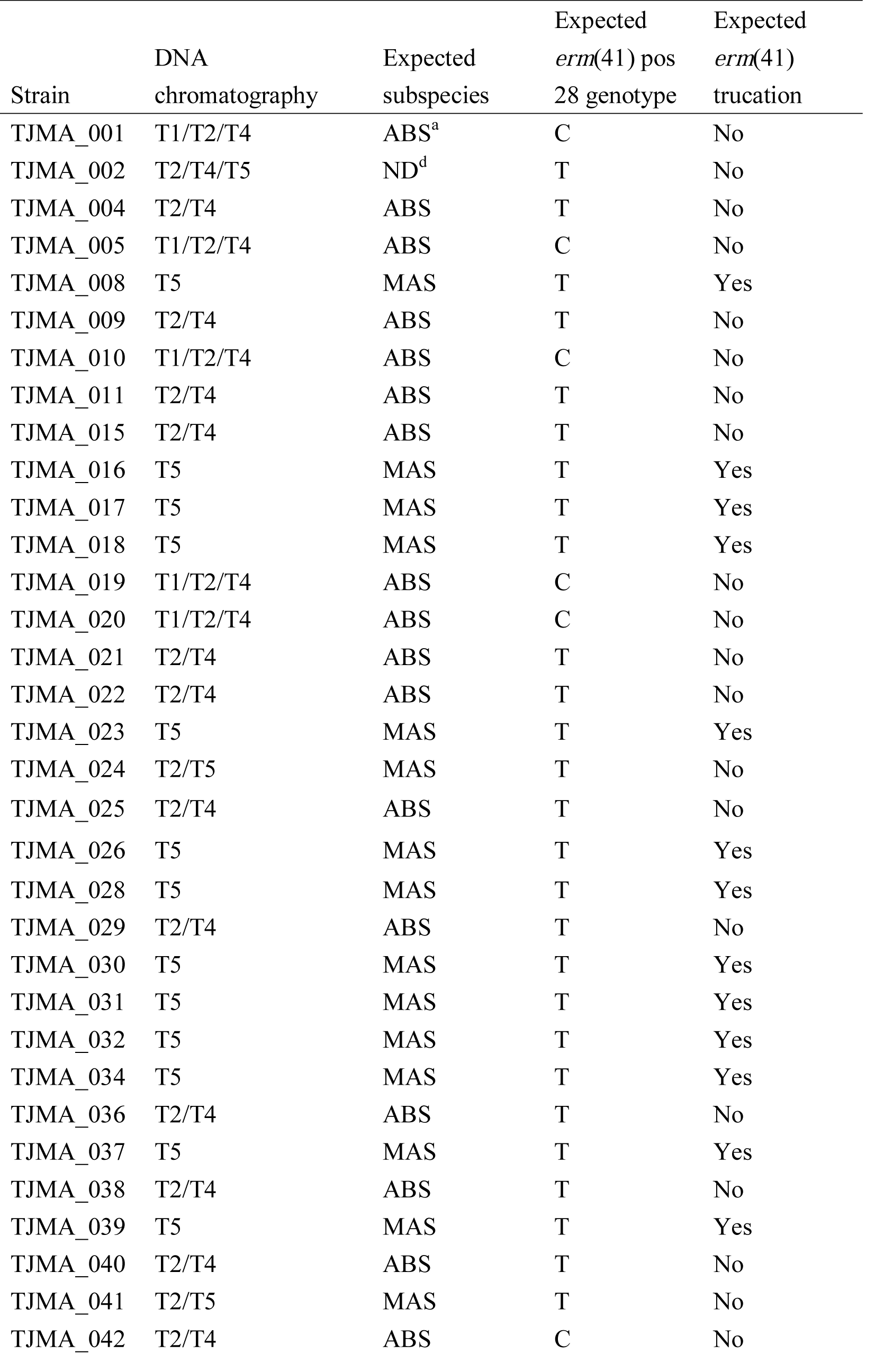
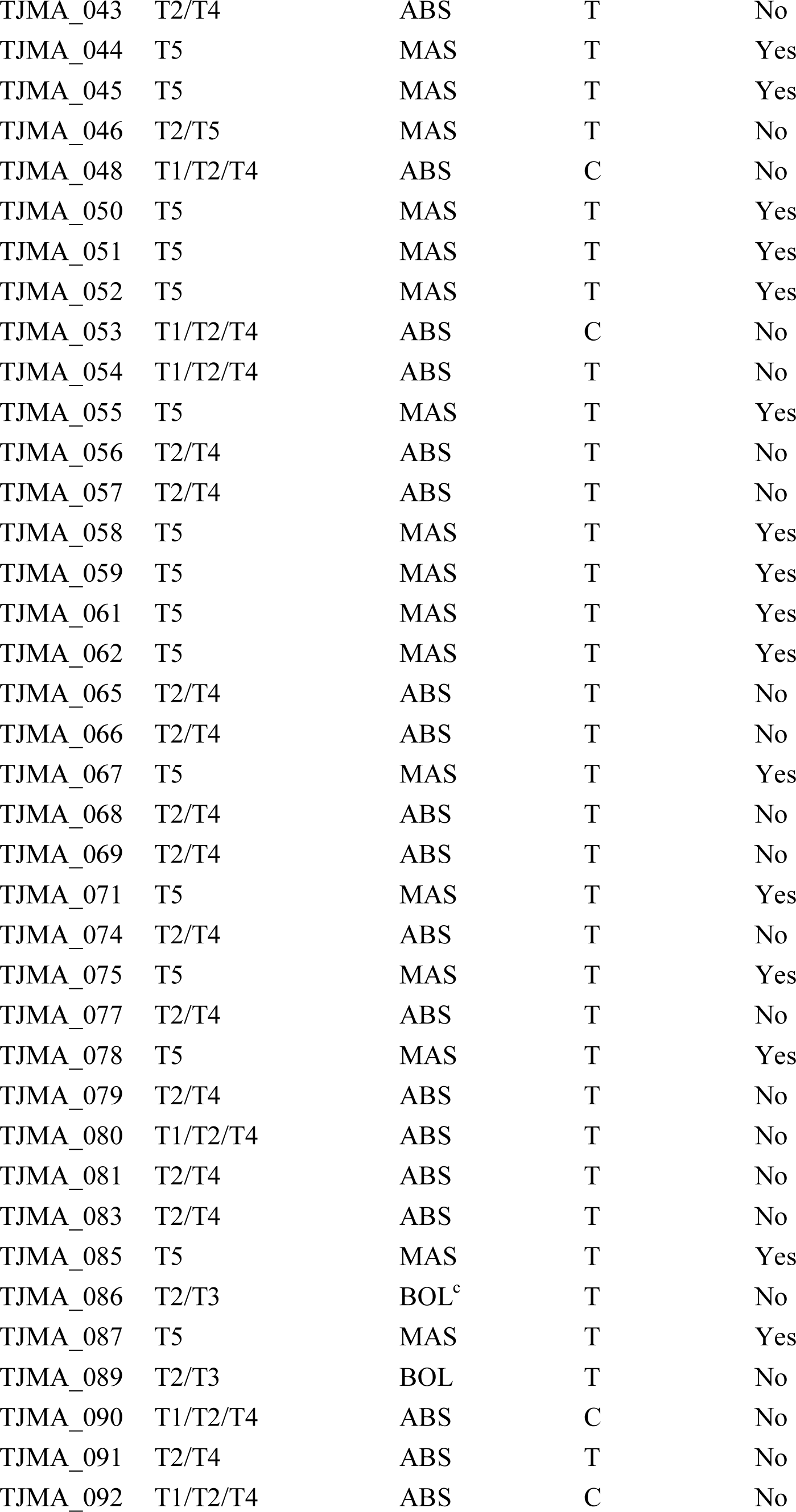
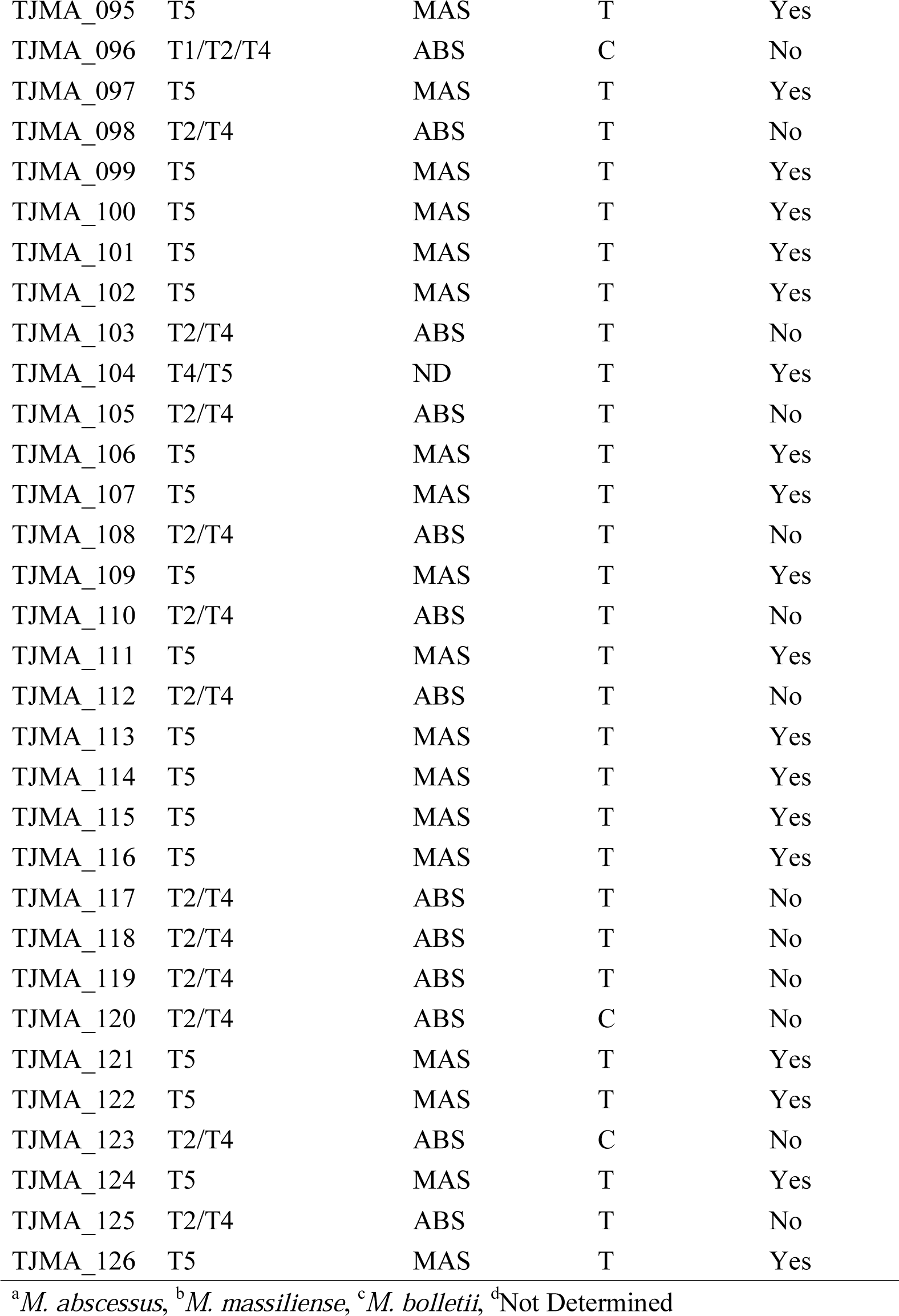
Subspecies and macrolide susceptibility identification of 103 clinical isolates from Taiwan using the DNA chromatography.

**Table S3.**
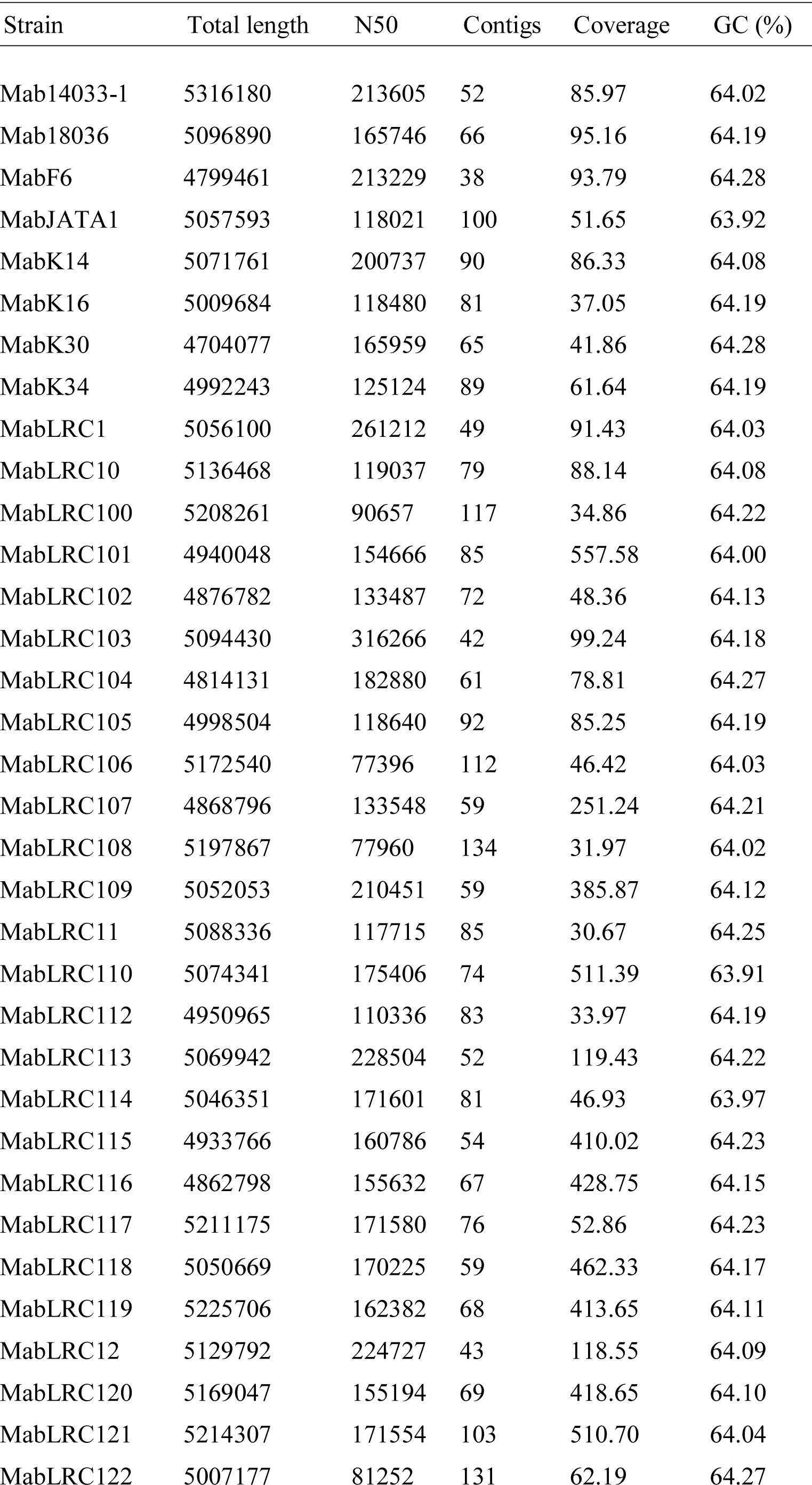
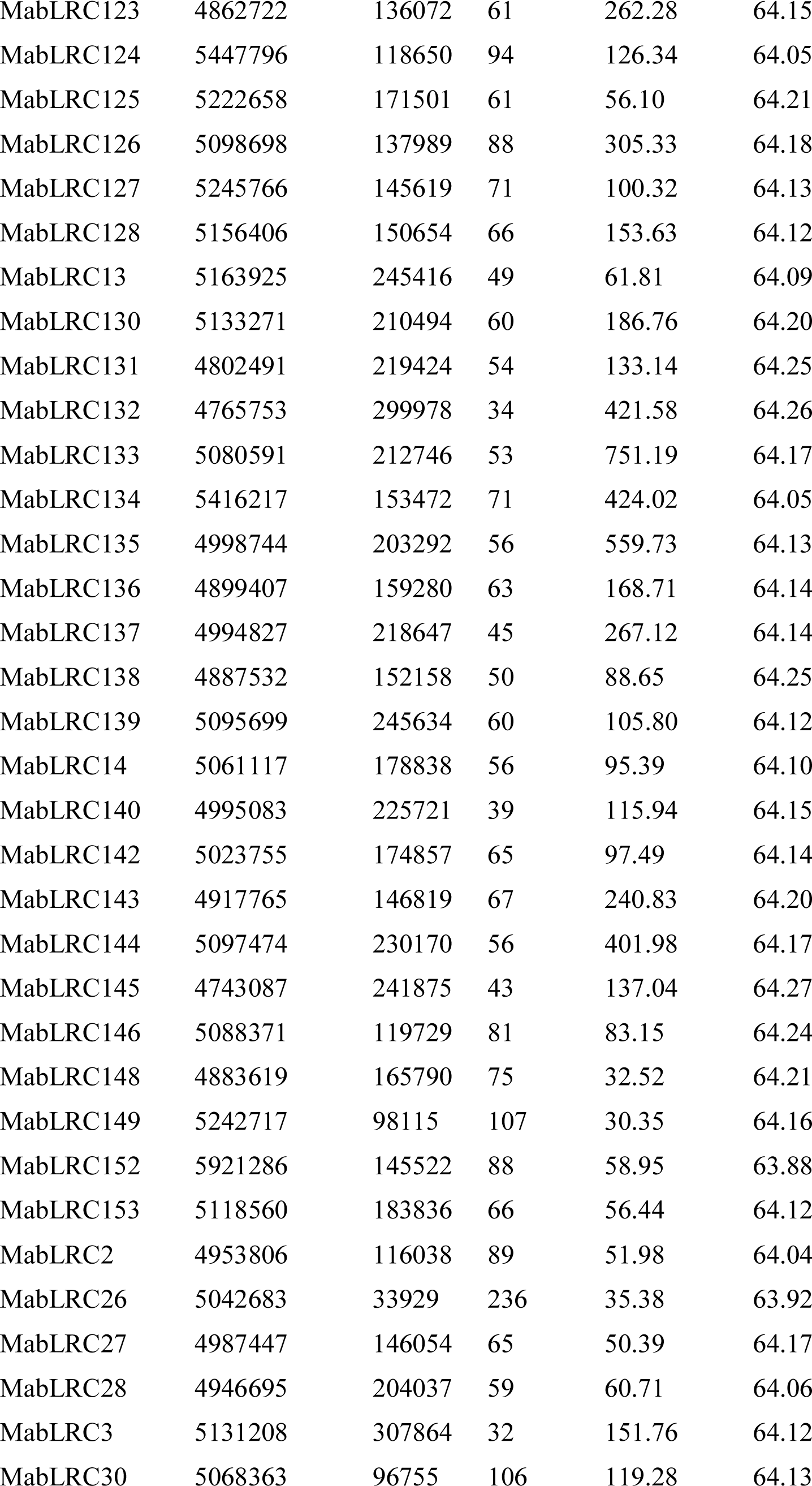
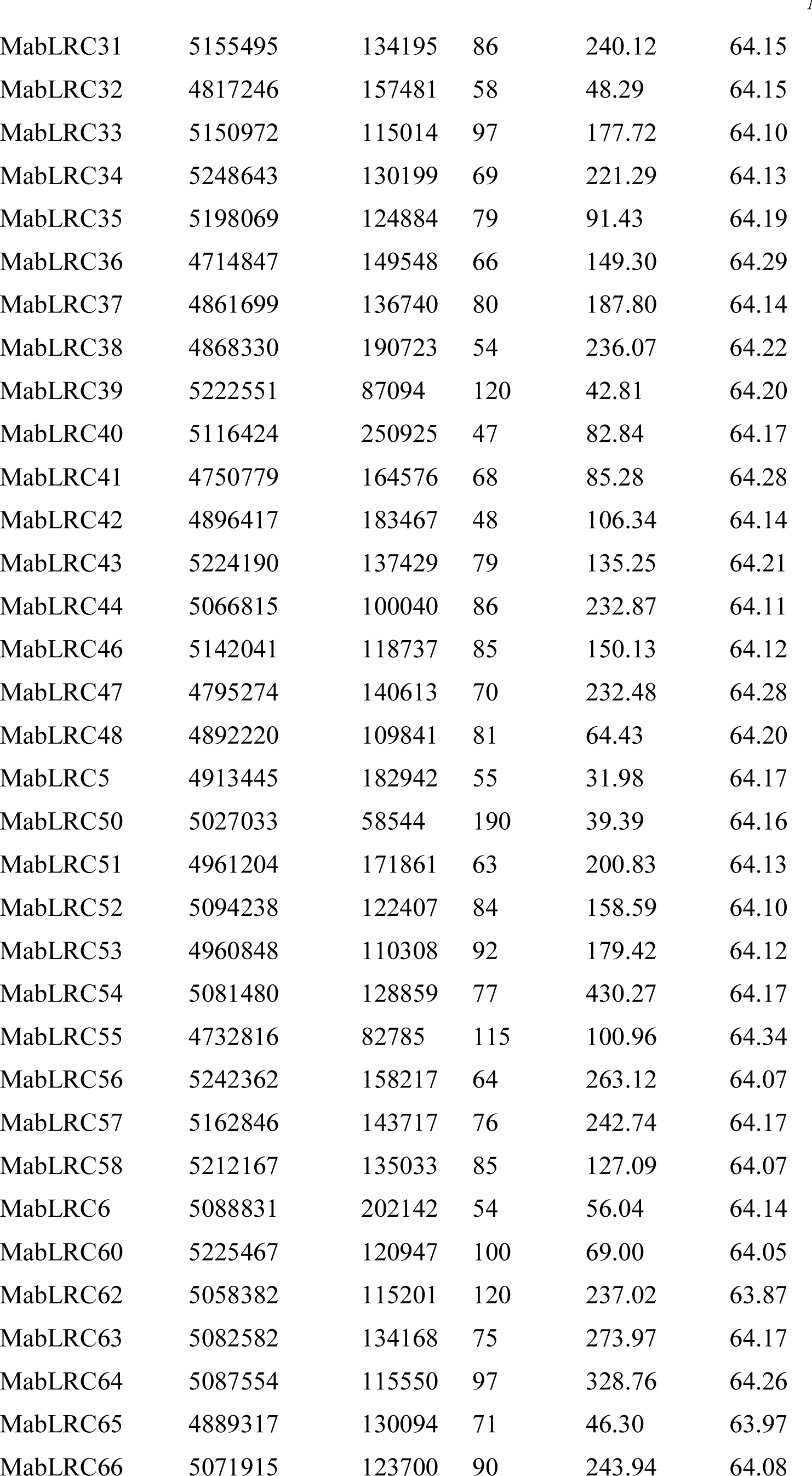
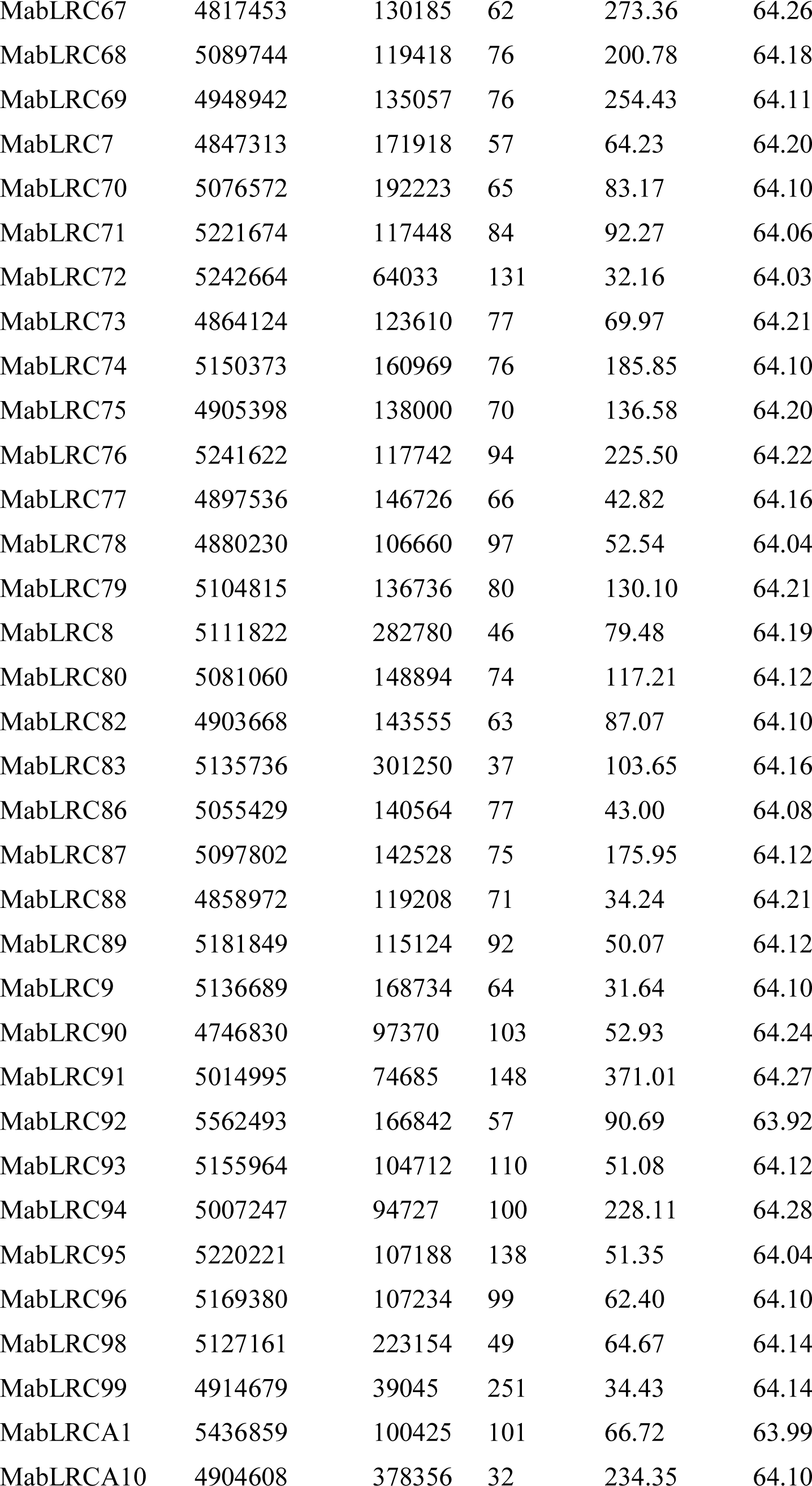
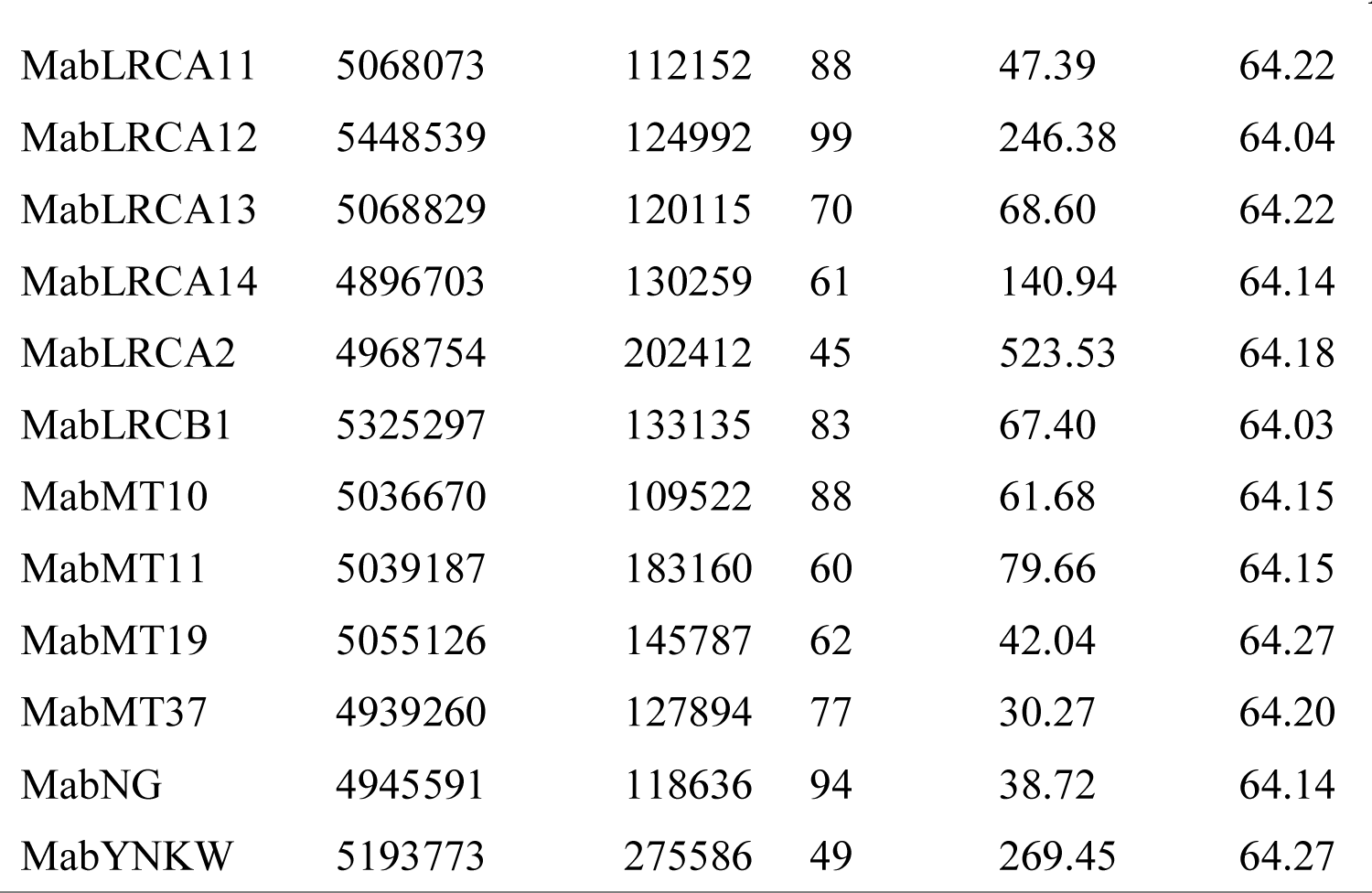
Basic statistics of WGS data of 148 environmental or clinical isolates from Japan

**Table S4.**
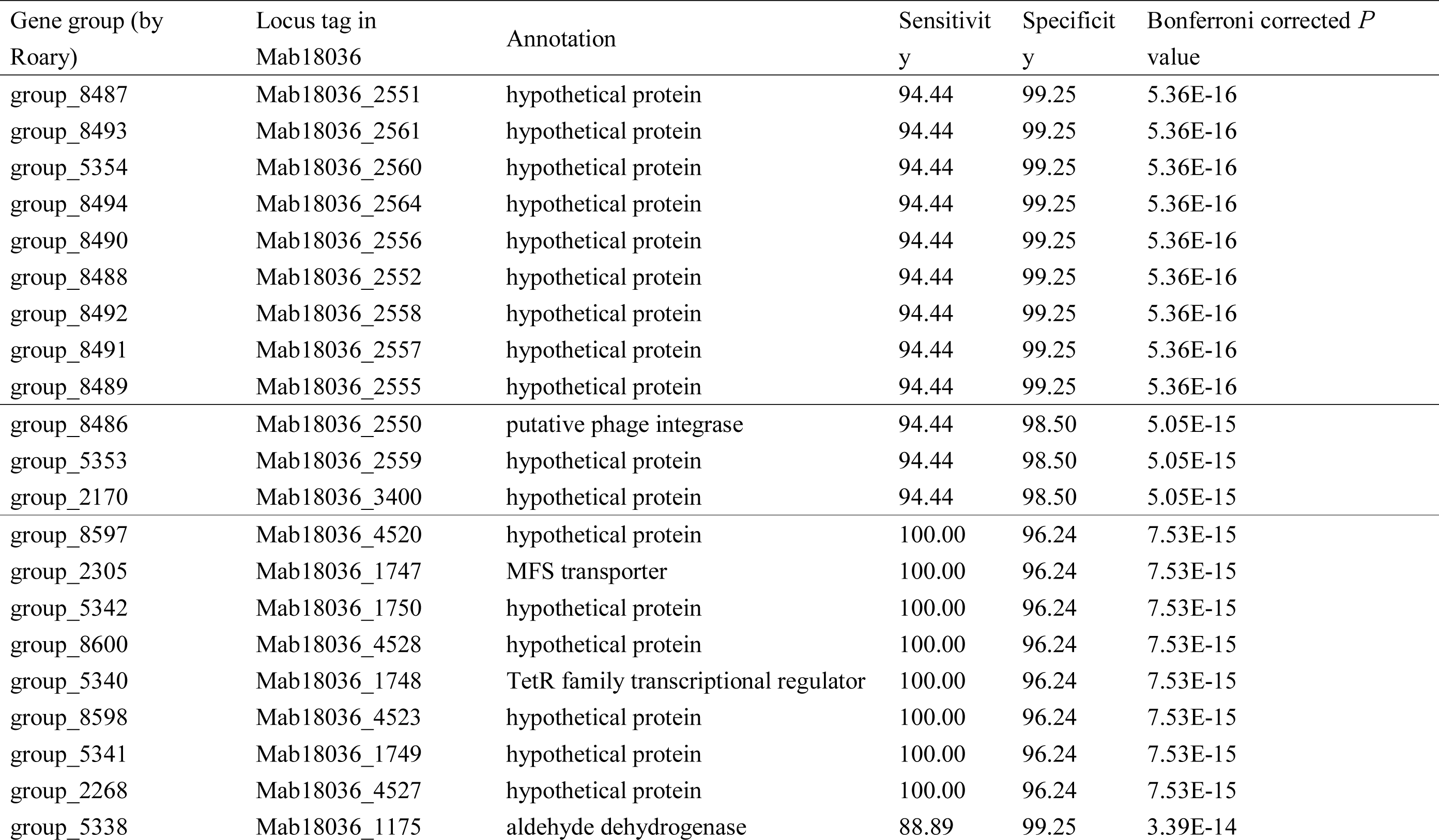
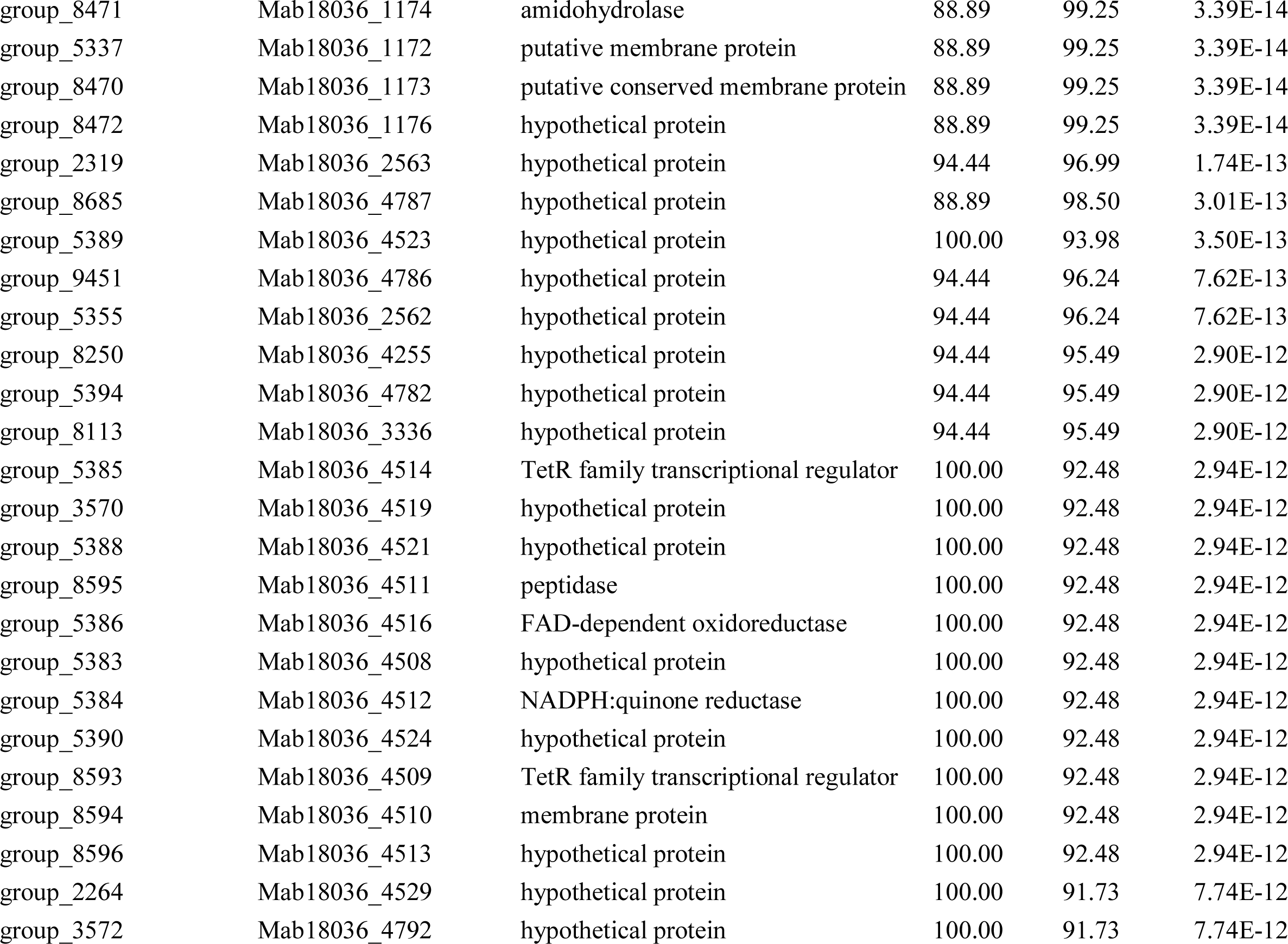
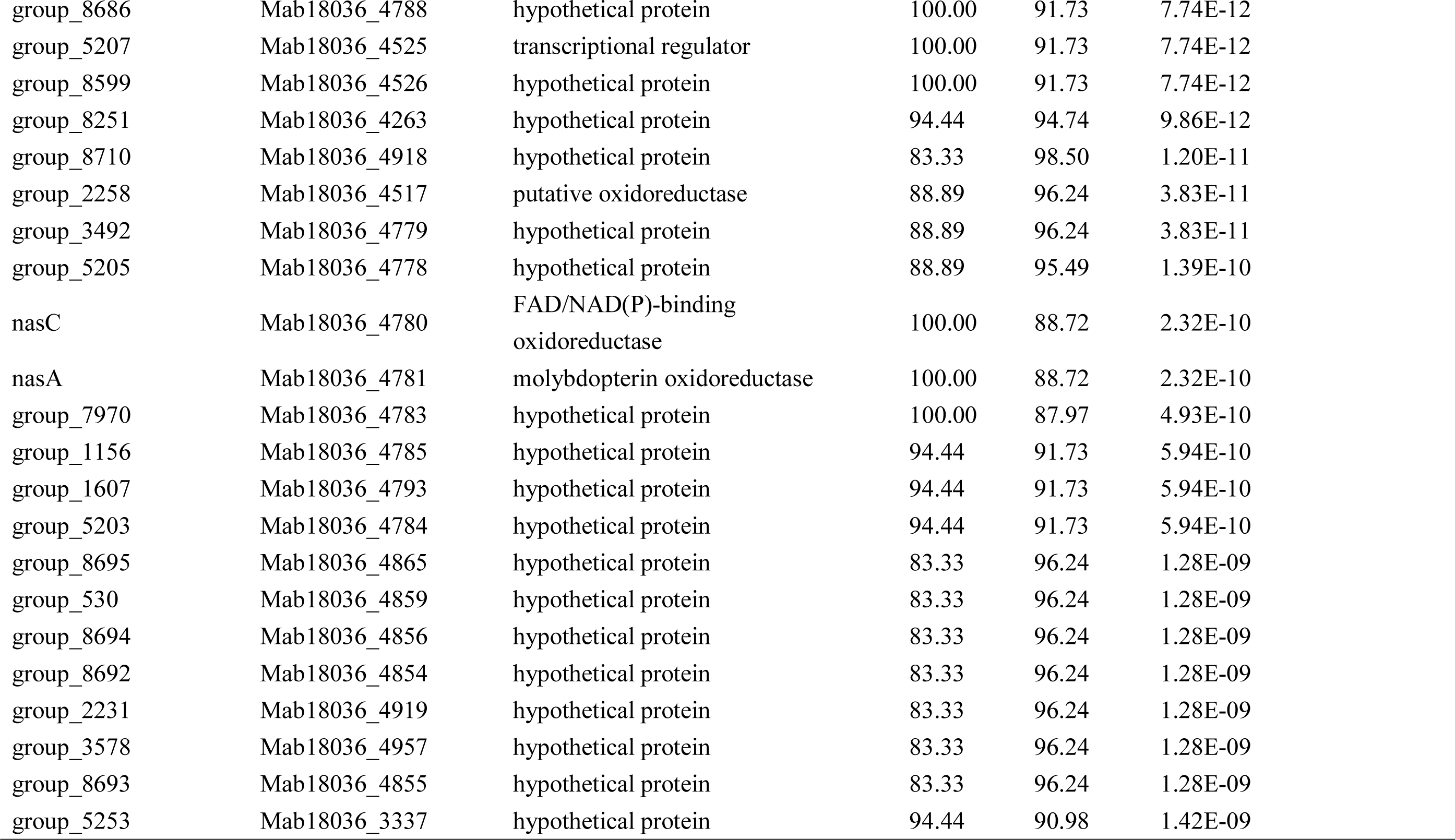
Genes associated with the lineage to which all *M. abscessus* with the *erm*(41) T28C sequevar belonged.

| Gene group (by Roary) | Locus tag in Mab18036 | Annotation | Sensitivity | Specificity | Bonferroni corrected *P* value |
| --- | --- | --- | --- | --- | --- |
| group_8487 | Mab18036_2551 | hypothetical protein | 94.44 | 99.25 | 5.36E-16 |
| group_8493 | Mab18036_2561 | hypothetical protein | 94.44 | 99.25 | 5.36E-16 |
| group_5354 | Mab18036_2560 | hypothetical protein | 94.44 | 99.25 | 5.36E-16 |
| group_8494 | Mab18036_2564 | hypothetical protein | 94.44 | 99.25 | 5.36E-16 |
| group_8490 | Mab18036_2556 | hypothetical protein | 94.44 | 99.25 | 5.36E-16 |
| group_8488 | Mab18036_2552 | hypothetical protein | 94.44 | 99.25 | 5.36E-16 |
| group_8492 | Mab18036_2558 | hypothetical protein | 94.44 | 99.25 | 5.36E-16 |
| group_8491 | Mab18036_2557 | hypothetical protein | 94.44 | 99.25 | 5.36E-16 |
| group_8489 | Mab18036_2555 | hypothetical protein | 94.44 | 99.25 | 5.36E-16 |
| group_8486 | Mab18036_2550 | putative phage integrase | 94.44 | 98.50 | 5.05E-15 |
| group_5353 | Mab18036_2559 | hypothetical protein | 94.44 | 98.50 | 5.05E-15 |
| group_2170 | Mab18036_3400 | hypothetical protein | 94.44 | 98.50 | 5.05E-15 |
| group_8597 | Mab18036_4520 | hypothetical protein | 100.00 | 96.24 | 7.53E-15 |
| group_2305 | Mab18036_1747 | MFS transporter | 100.00 | 96.24 | 7.53E-15 |
| group_5342 | Mab18036_1750 | hypothetical protein | 100.00 | 96.24 | 7.53E-15 |
| group_8600 | Mab18036_4528 | hypothetical protein | 100.00 | 96.24 | 7.53E-15 |
| group_5340 | Mab18036_1748 | TetR family transcriptional regulator | 100.00 | 96.24 | 7.53E-15 |
| group_8598 | Mab18036_4523 | hypothetical protein | 100.00 | 96.24 | 7.53E-15 |
| group_5341 | Mab18036_1749 | hypothetical protein | 100.00 | 96.24 | 7.53E-15 |
| group_2268 | Mab18036_4527 | hypothetical protein | 100.00 | 96.24 | 7.53E-15 |
| group_5338 | Mab18036_1175 | aldehyde dehydrogenase | 88.89 | 99.25 | 3.39E-14 |
| group_8471 | Mab18036_1174 | amidohydrolase | 88.89 | 99.25 | 3.39E-14 |
| group_5337 | Mab18036_1172 | putative membrane protein | 88.89 | 99.25 | 3.39E-14 |
| group_8470 | Mab18036_1173 | putative conserved membrane protein | 88.89 | 99.25 | 3.39E-14 |
| group_8472 | Mab18036_1176 | hypothetical protein | 88.89 | 99.25 | 3.39E-14 |
| group_2319 | Mab18036_2563 | hypothetical protein | 94.44 | 96.99 | 1.74E-13 |
| group_8685 | Mab18036_4787 | hypothetical protein | 88.89 | 98.50 | 3.01E-13 |
| group_5389 | Mab18036_4523 | hypothetical protein | 100.00 | 93.98 | 3.50E-13 |
| group_9451 | Mab18036_4786 | hypothetical protein | 94.44 | 96.24 | 7.62E-13 |
| group_5355 | Mab18036_2562 | hypothetical protein | 94.44 | 96.24 | 7.62E-13 |
| group_8250 | Mab18036_4255 | hypothetical protein | 94.44 | 95.49 | 2.90E-12 |
| group_5394 | Mab18036_4782 | hypothetical protein | 94.44 | 95.49 | 2.90E-12 |
| group_8113 | Mab18036_3336 | hypothetical protein | 94.44 | 95.49 | 2.90E-12 |
| group_5385 | Mab18036_4514 | TetR family transcriptional regulator | 100.00 | 92.48 | 2.94E-12 |
| group_3570 | Mab18036_4519 | hypothetical protein | 100.00 | 92.48 | 2.94E-12 |
| group_5388 | Mab18036_4521 | hypothetical protein | 100.00 | 92.48 | 2.94E-12 |
| group_8595 | Mab18036_4511 | peptidase | 100.00 | 92.48 | 2.94E-12 |
| group_5386 | Mab18036_4516 | FAD-dependent oxidoreductase | 100.00 | 92.48 | 2.94E-12 |
| group_5383 | Mab18036_4508 | hypothetical protein | 100.00 | 92.48 | 2.94E-12 |
| group_5384 | Mab18036_4512 | NADPH:quinone reductase | 100.00 | 92.48 | 2.94E-12 |
| group_5390 | Mab18036_4524 | hypothetical protein | 100.00 | 92.48 | 2.94E-12 |
| group_8593 | Mab18036_4509 | TetR family transcriptional regulator | 100.00 | 92.48 | 2.94E-12 |
| group_8594 | Mab18036_4510 | membrane protein | 100.00 | 92.48 | 2.94E-12 |
| group_8596 | Mab18036_4513 | hypothetical protein | 100.00 | 92.48 | 2.94E-12 |
| group_2264 | Mab18036_4529 | hypothetical protein | 100.00 | 91.73 | 7.74E-12 |
| group_3572 | Mab18036_4792 | hypothetical protein | 100.00 | 91.73 | 7.74E-12 |
| group_8686 | Mab18036_4788 | hypothetical protein | 100.00 | 91.73 | 7.74E-12 |
| group_5207 | Mab18036_4525 | transcriptional regulator | 100.00 | 91.73 | 7.74E-12 |
| group_8599 | Mab18036_4526 | hypothetical protein | 100.00 | 91.73 | 7.74E-12 |
| group_8251 | Mab18036_4263 | hypothetical protein | 94.44 | 94.74 | 9.86E-12 |
| group_8710 | Mab18036_4918 | hypothetical protein | 83.33 | 98.50 | 1.20E-11 |
| group_2258 | Mab18036_4517 | putative oxidoreductase | 88.89 | 96.24 | 3.83E-11 |
| group_3492 | Mab18036_4779 | hypothetical protein | 88.89 | 96.24 | 3.83E-11 |
| group_5205 | Mab18036_4778 | hypothetical protein | 88.89 | 95.49 | 1.39E-10 |
| nasC | Mab18036_4780 | FAD/NAD(P)-binding<br>oxidoreductase | 100.00 | 88.72 | 2.32E-10 |
| nasA | Mab18036_4781 | molybdopterin oxidoreductase | 100.00 | 88.72 | 2.32E-10 |
| group_7970 | Mab18036_4783 | hypothetical protein | 100.00 | 87.97 | 4.93E-10 |
| group_1156 | Mab18036_4785 | hypothetical protein | 94.44 | 91.73 | 5.94E-10 |
| group_1607 | Mab18036_4793 | hypothetical protein | 94.44 | 91.73 | 5.94E-10 |
| group_5203 | Mab18036_4784 | hypothetical protein | 94.44 | 91.73 | 5.94E-10 |
| group_8695 | Mab18036_4865 | hypothetical protein | 83.33 | 96.24 | 1.28E-09 |
| group_530 | Mab18036_4859 | hypothetical protein | 83.33 | 96.24 | 1.28E-09 |
| group_8694 | Mab18036_4856 | hypothetical protein | 83.33 | 96.24 | 1.28E-09 |
| group_8692 | Mab18036_4854 | hypothetical protein | 83.33 | 96.24 | 1.28E-09 |
| group_2231 | Mab18036_4919 | hypothetical protein | 83.33 | 96.24 | 1.28E-09 |
| group_3578 | Mab18036_4957 | hypothetical protein | 83.33 | 96.24 | 1.28E-09 |
| group_8693 | Mab18036_4855 | hypothetical protein | 83.33 | 96.24 | 1.28E-09 |
| group_5253 | Mab18036_3337 | hypothetical protein | 94.44 | 90.98 | 1.42E-09 |

